# HNRNPH1 destabilizes the G-quadruplex structures formed by G-rich RNA sequences that regulate the alternative splicing of an oncogenic fusion transcript

**DOI:** 10.1101/2022.04.18.488656

**Authors:** Tam Vo, Tayvia Brownmiller, Katherine Hall, Tamara L. Jones, Sulbha Choudhari, Ioannis Grammatikakis, Katelyn R. Ludwig, Natasha J. Caplen

## Abstract

In the presence of physiological monovalent cations, thousands of RNA G-rich sequences can form parallel G-quadruplexes (G4s) unless RNA-binding proteins inhibit, destabilize, or resolve the formation of such secondary RNA structures. Here, we have used a disease-relevant model system to investigate the biophysical properties of the RNA-binding protein HNRNPH1’s interaction with G-rich sequences. We demonstrate the importance of two *EWSR1*-exon 8 G-rich regions in mediating the exclusion of this exon from the oncogenic *EWS-FLI1* transcripts expressed in a subset of Ewing sarcomas, using complementary analysis of tumor data, long-read sequencing, and minigene studies. We determined that HNRNPH1 binds the *EWSR1*-exon 8 G-rich sequences with low nM affinities irrespective of whether in a non-G4 or G4 state but exhibits different kinetics depending on RNA structure. Specifically, HNRNPH1 associates and dissociates from G4-folded RNA faster than the identical sequences in a non-G4 state. Importantly, we demonstrate using gel shift and spectroscopic assays that HNRNPH1, particularly the qRRM1-qRRM2 domains, destabilizes the G4s formed by the *EWSR1*-exon 8 G-rich sequences in a non-catalytic fashion. Our results indicate that HNRNPH1’s binding of G-rich sequences favors the accumulation of RNA in a non-G4 state and that this contributes to its regulation of RNA processing.

## INTRODUCTION

The inclusion or exclusion of specific sequences during mRNA biogenesis is a fundamental mechanism that expands the human transcriptome’s complexity (1–3). Critical elements defining a specific transcript variant’s expression include *cis*-acting sequences, which, depending on their position within exons or introns, act as splicing enhancers or suppressors, the trans-acting RNA-binding proteins (RBPs) that bind these sequences, and the local and long-range intramolecular interactions that induce RNA folding (4–6). Members of the heterogeneous nuclear RNA proteins (hnRNPs) are essential regulators of RNA processing, including alternative splicing (7, 8). The primary oncogenic mutation in most cases of the pediatric tumor Ewing sarcoma involves chromosome 22 translocations in which the breakpoint occurs in either intron 7 or intron 8 of the *EWSR1* gene (**Figure 1A**). We identified one of the members of the HNRNPH/F subfamily of hnRNPs, HNRNPH1, as critical for excluding *EWSR1*-exon 8 from the fusion pre-mRNAs expressed in the subset of Ewing sarcomas with *EWSR1*-intron 8 breakpoints (9). Failure to exclude *EWSR1*-exon 8 results in an mRNA that includes a premature stop codon, leading to loss of the expression of the EWS-FLI1 oncoprotein that Ewing sarcoma cells need for survival (9) (**Figure 1A**).

**Figure 1:**
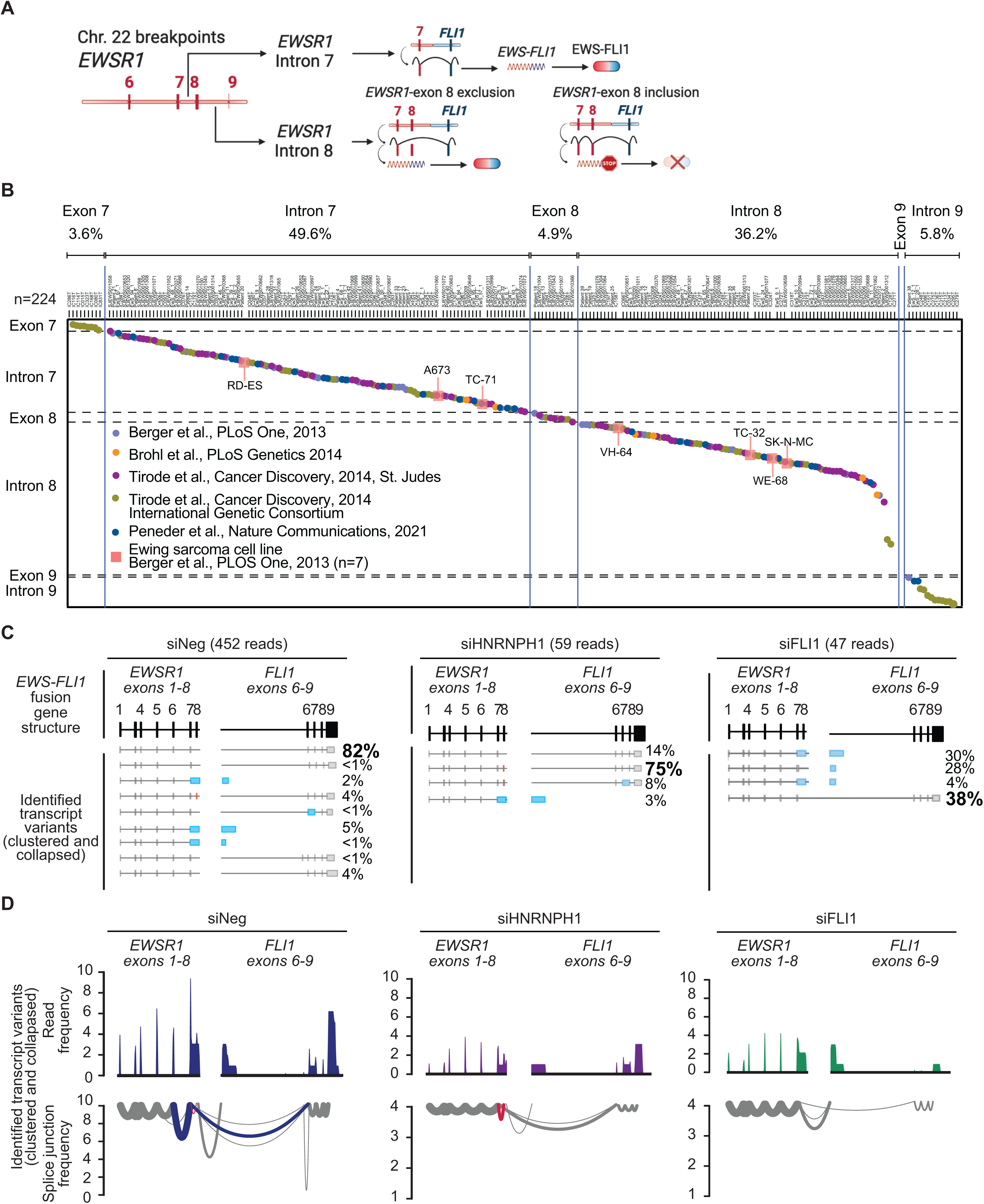
The HNRNPH1-dependent processing of the *EWS-FLI1* transcript expressed in an over 35% of Ewing sarcomas. **A.** Schematic of the effect of Chromosome 22 breakpoint positions on the processing *EWS-FLI1* transcripts and the expression of the EWS-FLI1 fusion oncoprotein. **B.** A meta-analysis of *EWSR1* breakpoints in a sample of 224 EWS-FLI1-driven tumors and 7 Ewing sarcoma cell lines. We extracted these results from Berger *et al*., 2013 (51), Tirode et al., 2014 (International Cancer Genome Consortium and St. Jude’s Cloud PeCan portal (https://pecan.stjude.cloud/) (52), Brohl *et al*., 2014 (53) and Peneder *et al*., 2021 (54). **C.** *EWS-FLI1* transcripts (clustered and collapsed) detected by long-read RNA sequences of analysis of TC-32 cells transfected with the indicated siRNAs and the percentage of reads mapping to each transcript variant. The top track indicates the consensus structure of *EWSR1* and *FLI1* intron and exon sequences, the bottom track, schematics of each detected transcript variant (clustered and collapsed). Red rectangle indicates *EWSR1*-exon 8. Light blue rectangles indicate retained intronic sequences. **D.** Long-read sequencing of *EWS-FLI1* transcripts expressed in TC-32 cells transfected with the indicated siRNAs. The top track shows the transcript variant mapping frequency. The bottom track shows Sashimi plots. Blue lines indicate transcripts excluding *EWSR1*-exon 8. Red lines indicate transcripts including *EWSR1*-exon 8. **A** created in BioRender. **C** and **D** generated using Gviz.

HNRNPH1 binds guanine (G)-rich sequences, and its binding of G-tracts within intronic or exonic sequences, in combination with 5’ splice sites of varying strengths, is a determinant of exon inclusion or exclusion (10, 11). Several recent studies have highlighted the interaction of HNRNPH1 and other members of the HNRNPH/F protein family with G-rich RNA sequences that, at physiologically relevant monovalent salt concentrations (e.g., K^+^ or Na^+^), could form G-quadruplex (G4) structures (11–14). G4 structures comprise stacks of two or more planer G-tetrads formed via Hoosgteen-hydrogen bonds. Transcriptome-wide *in vitro* profiling methods using reverse transcriptase stalling and next-generation sequencing has uncovered thousands of RNA G4s (rG4s), most of which adopt parallel structures fitting the consensus G_≥2_N_1-7_G_≥2_N_1-7_ G_≥2_N_1-7_G_≥2_ motif that stack two or three guanine-tetrads in a regular structure (15, 16). Of note, the additional 2’OH group in the ribose ring and C3’-endo pucker conformations, differences in water activities during quadruplex formation, and increased intramolecular hydrogen bonding are all features that promote rG4s to form a parallel topology that is more stable than that formed by DNA G4s (17–20). Canonical G4 RNA structures consist of three stacked guanine-tetrads, but recent studies have indicated that G-rich RNAs can form more complex structures, including the formation of non-canonical three guanine-tetrads with a less regular looping or bulges or a stack of two guanine-tetrads forming a two-tier quartet structure (15,21,22). However, it remains unclear whether these structures form within cells. For example, Guo and Bartel proposed that in mammalian cells, G-tract binding proteins such as members of the HNRNPH/F subfamily of hnRNPs or SRSF1/2 inhibit the formation of rG4 structures (16). Recently, we demonstrated that HNRNPH1 binds at least one G-rich sequence present at the 3’ end of *EWSR1*-exon 8 using RNA oligomers and that in the presence of cations, this RNA folds into a G4 structure (23). We also observed that Ewing sarcoma cell lines dependent on the exclusion of *EWSR1*-exon 8 to express the fusion oncoprotein EWS-FLI1 exhibit loss of cell viability following exposure to the pan-G4-binder pyridostatin at lower concentrations than other cell lines (23).

In addition to our studies implicating HNRNPH1 in processing the oncogenic fusion transcripts expressed in a subset of Ewing sarcomas, several recent reports have described disease-associated sequence changes in *HNRNPH/F* genes. For example, multiple reports have linked mutations in *HNRNPH1* or *HNRNPH2* to rare neurodevelopmental syndromes that exhibit variations in phenotype but include dysmorphic features and intellectual disability (24–30). The described mutations include missense mutations, frameshift and in-frame deletions, and partial gene duplications (30, 31). In addition, a study of driver genes in ulcerative colitis linked an increased risk of colorectal cancer to somatic mutations in HNRNPF (32), and recent studies of mantle-cell lymphoma samples have identified mutations in HNRNPH1 at an increased frequency (33, 34). There are also reports of disease-associated changes in sequences bound by HNRNPF/H proteins, as illustrated by HNRNPH1’s regulation of the expression of *OPRM1* transcript variants. *OPRM1* codes for the mu-opioid receptor and a single nucleotide (nt) polymorphism (G-to-A) in an intron splice enhancer sequence bound by HNRNPH1 modulates protein binding, increasing the likelihood of a specific exon exclusion event (35). Related to this observation are the mouse-based genetic studies that have linked Hnrnph1 function to phenotypes associated with addictive behaviors (36–40). However, to understand the effect of HNRNPF/H mutations or variations in the sequences bound by these proteins, we need to further study their interaction with specific sequences and analyze the effect on the RNAs they bind.

The HNRNPH1/H2/F proteins interact with G-rich sequences through quasi-RNA recognition motif (qRRM) domains that lack the conserved aromatic residues observed in RBPs (41, 42). Studies of the HNRNPF protein show that the qRRM domains form a series of *β*_1_/*α*_1_, *β*_2_/*β*_3_, and *α*_2_/*β*_4_ loops in the presence of RNA (AGGGAU) that is typical of most RRM domains, but qRRMs interact with RNA via these loops rather than through interactions with *β*-sheet surfaces (43). X-ray-crystallography and NMR studies show that the qRRM1-qRRM2 domains of HNRNPH1/H2 switch between a compact state, resulting in RNA recognition by both qRRMs of a single G-rich sequence and an extended state where it can bind to two different G-tracts. HNRNPH1/H2 primarily form compact states, and the linker between the two qRRMs remains rigid, differing from HNRNPF, which exhibits a flexible and extended state (44). Consistent with the assertion that the binding of HNRNPH/F proteins inhibits rG4 formation (16), Dominguez and co-workers showed that a single qRRM domain of HNRNPF could alter the splicing of the *BCL-X* pre-RNA by inhibiting the formation of the intramolecular bonds required for the formation of a secondary structure (43). Furthermore, a study by Samatanga and colleagues also showed that qRRM3 of HNRNPF binds exclusively to unfolded G-tracts and not to rG4s and that the protein-RNA complex forms before G4 formation (45).

Here, we have used the HNRNPH1-dependent exclusion of *EWSR1*-exon 8 from *EWS-FLI1* transcripts as a model system to investigate HNRNPH1’s interaction with G-rich sequences. We demonstrate the relevance of the model system using tumor data and cell-based analysis, highlighting that though the HNRNPF and HNRNPH1/H2 proteins are highly homologous (70% to over 90% homology), only HNRNPH1 is responsible for the alternative splicing of *EWSR1*-exon 8 in the context of the *EWS-FLI1* transcript containing this exon. Using a minigene system, we determine that G-tracts in the 3’ end of the exon are critical determinants of *EWSR1*-exon 8 splicing. Next, using RNA oligomers corresponding to these G-rich regions, we demonstrate that RNA transcripts containing these sequences would form G4 structures under physiologically relevant salt conditions. We predict these rG4 structures will contain guanine-tetrads consisting of two-tier quartets, a motif enriched in exonic sequences (23), and that may have thermodynamic properties compatible with regulatory functions (46) unless a protein inhibits RNA folding or a protein destabilizes or resolves the folded RNA. To address whether HNRNPH1 can alter the propensity of the *EWSR1*-exon 8 G-rich sequences to form these two-tier quartet structures, we used orthogonal Fluorescence Resonance Energy Transfer (FRET) and Circular Dichroism (CD) assays to establish that HNRNPH1 can alter the structural conformation of RNA in a G4 state. Furthermore, analysis of protein-RNA interactions using biosensor Bio-layer Interferometry (BLI) analysis suggests that the kinetics of HNRNPH1’s interaction with the G-rich sequences in *EWSR1*-exon 8 favors the accumulation of these RNA regions in a non-G4 state. In brief, the association kinetics of HNRNPH1’s binding of G-rich RNA unable to form a G4-state favors the maintenance of this state. At the same time, the association kinetics of HNRNPH1’s binding and subsequent destabilization of G4-RNAs promotes a shift from a G4 to a non-G4 state. Our results further our understanding of how a member of the HNRNPHF family of proteins can contribute to the rapid changes in RNA structure needed to regulate RNA processing and how their binding of the *cis*-regulatory sequences that have the potential to form complex secondary structures may represent a critical aspect of their functions as alternative splicing factors.

## MATERIAL AND METHODS

### Reagents

HNRNPH1 full-length protein was purchased from OriGene Technologies (Rockville, MD). The recombinant monoclonal antibody against DNA/RNA Quadruplex structures, BG4, was purchased from Absolute Antibody (Boston, MA). The expression and purification of the HNRNPH1 truncated proteins qRRM1-2 and qRRM2-3 (Creative BioMart, Shirley, NY) were performed as described previously (23). Synthetic biotinylated RNAs: rG1, 5’-Bio/rArCrCrGrGrGrGrCrArGrGrGrGrArArGrArGrGrGrGrGrArUrU-3’; rG2, 5’-Bio/rGrGrUrGrGrGrCrGrGrGrGrArGrGrArGrGrArCrGrCrGrGrUrGrGrArArU-3’; rG3, 5’- Bio/rGrGrGrCrGrGrGrGrArGrGrArGrGrArCrGrCrGrGrUrGrGrArArUrGrGrG-3’), MALAT1, 5’- rArCrGrGrUrUrGrGrGrArUrUrGrGrUrGrGrGrGrUrGrGrGrUrU-3’/Bio; rG1-FRET 5’- Cy5/rArCrCrGrGrGrGrCrArGrGrGrGrArArGrArGrGrGrGrGrArUrU/Cy3-3’; and rG3-FRET 5’- Cy5/rGrGrGrCrGrGrGrGrArGrGrArGrGrArCrGrCrGrGrUrGrGrArArUrGrGrG/Cy3-3’ were purchased from Integrated DNA Technology (IDT, Coralville, IA). We also purchased the rG1mt and rG1-7dG and the rG3-mt and rG3-7dG RNA oligomers from IDT. See **Figures 4** and **S3** for the sequences of these RNA oligomers.

The RNA oligomers were desalted using the Slide-A-Lyzer™ MINI Dialysis Device (catalog number 88403) from Thermo Fisher Scientific (Waltham, MA) in double-distilled water (ddH2O) following the manufacturer’s instructions. Briefly, 200 µL of 100 µM of each RNA sample was loaded into the dialysis device and dialyzed the RNA in a 15 mL conical tube containing 14 mL ddH2O on an orbital shaker for 4 hours at room temperature and then at 4°C overnight. Unless otherwise stated, RNA oligomers were resuspended in a 10 mM Tris-HCl pH 7.4 and 1 mM EDTA buffer only or plus KCl to reach the required salt concentration per sample – 1 to 150 mM KCl. We heat-denatured the RNA oligomers at 95°C for at 2 to 5 minutes (mins), followed by cooling to room temperature for a minimum of 30 mins and a maximum of 90 mins prior to experimentation.

The pRHCGlo plasmid was obtained from Addgene (Watertown, MA; Catalog number, 80169) (47). For RNAi studies, siRNAs were purchased from Qiagen (Germantown, MD) or Thermo Fisher Scientific (Ambion) as follows: siNeg (Qiagen, SI03650318), siHNRNPH1 (Ambion, s6730, GGAUUUGGGUCAGAUAGAUtt), siHNRNPH2 (Ambion, 145362, GCACUAAAUAGCUACUCCAtt), siHNRNPF, (Qiagen, SI00300461, AAGCGUUCGUGCAGUUUGCCU), and siFLI1 (Ambion s5266, CAAACGAUCAGUAAGAAUAtt). Custom PCR primers (DNA) were purchased from Thermo Fisher Scientific: RSV5U (Forward) 5’-CATTCACCACATTGGTGTGC-3’(47); RTRHC 5’-GGGCTTGCAGCAACAGTAAC-3’ (First-strand synthesis and Reverse primer) (47); *HNRNPH1*, Forward - 5’-TACACATGCGGGGATTACCTT-3’, Reverse - 5’-CTTCACCAGTTACTCTGCCATC-3’; *HNRNPH2*, Forward - 5’-CTCGCTATAGCCGTTTGAGG-3’, Reverse - 5’-GTGATTGTTGGGCTCTTGGT-3’; *HNRNPF,* Forward - 5’-GCCTGGTAGCAACAGAAACC-3’, Reverse - 5’-GTGATCTTGGGTGTGGC-3’; *EWS-FLI1*, *EWSR1*-exon 7, Forward – 5’-ATCCTACAGCCAAGCTCCAA-3’, Reverse *FLI1* exon 7 Reverse 5’ GGCCGTTGCTCTGTATTCTTAC-3’; *NACA* - Forward - 5’-ACAAGAGCCCTGCTTCAGAT-3’ Reverse – 5’-GCTGCTAGTTGTGCTTGCTG-3’.

### Biological Resources

TC-32 cells were obtained from the Pediatric Oncology Branch, CCR, NCI, and HEK-293T cells from ATCC (Manassas, Virginia). Short tandem repeat (STR) fingerprint information (ATCC) are available upon request. Cells were grown in RPMI 1640 (Invitrogen, Carlsbad, CA) with 10% FBS (Invitrogen) and Plasmocin (InvivoGen USA, San Diego, CA) and checked for mycoplasma contamination using the MycoAlert Plus system (Lonza, Walkersville, MD) regularly, which was not detected.

### Circular Dichroism (CD) spectroscopy

CD spectra were recorded using a Chirascan Q100 spectropolarimeter from Applied Photophysics, Inc. (Beverly, MA) at room temperature. In brief, 250 µL per sample was loaded into the well of a 96-micro well plate, and each sample was transferred into a 1 nm pathlength microcuvette using an autosampler. Three spectra per sample were obtained, recording data with a data interval of 1 scan/nm. Each dataset was normalized by subtracting the values obtained using a blank sample and plotting the average value of the three replica spectra per sample. The thermal melting studies used 3 µM RNA in 10 mM Tris-HCl pH 7.4, 1 mM EDTA, and 25 mM KCl and assessed the change in molar ellipticity from 220 to 350 nm with the increasing temperature (25°C to 95°C) at a rate of 0.5°C/min and plotted results in Prism (GraphPad, San Diego, CA). CD spectra of the truncated HNRNPH1 domains (qRRM1-2 or qRRM2-3) were recorded at a constant 3 µM in the experimental buffer. For protein titration experiments, we added an increasing concentration (0 to 12 µM) of each truncated protein, HNRNPH1 (qRMM1-2 or qRRM2-3) to 3 µM RNA, assessing RNA:protein ratios of 1:0, 1:1, 1:2, 1:3, and 1:4.

### Bio-Layer Interferometry

Bio-Layer Interferometry (BLI) is a biosensor-based method that detects the wavelength shift that occurs following interaction of an immobilized ligand on a sensor chip, in this case, an RNA oligomer, and a biomolecule of interest, in this case, HNRNPH1. For our studies we used the Octet Red96e system, Fortébio (Fremont, CA). We purchased the 96-wells plates (cat #651209, Greiner Bio-One, Monroe, NC). Unless otherwise stated, we used BLI buffer containing 10 mM Tris-HCl (pH 7.4), 150 mM KCl, 0.005% (v/v) P-20, 1 mM EDTA for all BLI experiments. To prepare the biotin-labeled RNA, we heat denatured 20 nM RNA oligomer in 200 µL BLI buffer and then allowed the samples to cool at room temperature for up to 90 minutes. To prepare protein, we serially diluted HNRNPH1 full-length protein to obtain proteins concentrations ranging from 1 nM to 20 nM (in triplicate) in BLI buffer. To prepare the streptavidin-coated biosensor chips, we equilibrated each chip in BLI buffer for 15 minutes and then immobilized the annealed biotin-labeled RNA on the sensor chip surface for 120 seconds. Next, we added BLI buffer for 200 seconds to establish a stable baseline reading for each sensor, and then we dipped the sensor chip into the wells containing the protein samples. We used an association time of 400 seconds and a dissociation time of 2000 seconds. To neutralize and regenerate the sensor chip, we exposed the surface to 1M KCl buffer for 60 seconds, followed by 120 seconds in BLI buffer. We performed each BLI experiment in triplicate and present a representative sensorgram in each case. The results of the replicate datasets were fitted to a 1:1 binding model for kinetic analysis using BLI Octet Data evaluation software. Dissociation binding equilibrium (K_D_) value was calculated:

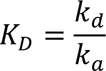

Where k_d_ is dissociation kinetic rate and k_a_ is association kinetic rate obtained from the fitting. We plotted the results in Prism (GraphPad).

### RNA pulldown

RNA pulldowns used the Pierce Magnetic RNA-Protein Pull-down Kit (Thermo Fisher Scientific). In brief, 50 µL of streptavidin magnetic beads were washed with 20 mM Tris and then incubated with 50 µL 1x RNA capture buffer and 50 pmol of each biotinylated RNA oligomer (20 mM Tris-HCl (pH 7.5) and 150 mM KCl). After washing with 20 mM Tris and 1x protein-RNA binding buffer, the RNA-bound beads were incubated with Master Mix, 10x protein-RNA binding buffer, 50% glycerol, and either 80 µg of whole cell lysate at 4°C for 30 minutes or 400 ng recombinant HNRNPH1 at room temperature for 1 hour. The protein-RNA-bound beads were washed with wash buffer, eluted, and the supernatant was collected for analysis by western blot. The eluted samples were heated at 95°C for 10 minutes with 1x reducing sample buffer and loaded into a 4%-20% tris-glycine gel (Thermo Fisher Scientific) at 30 µL/well. Blotting was performed using iBlot 2 Gel Transfer Device (Thermo Fisher Scientific). We detected endogenous HNRNPH1 using an anti-HNRNPH1 antibody (A300-511A, Bethyl Laboratories, Montgomery, TX) and an anti-rabbit HRP secondary antibody (7074, Cell Signaling Technology, Danvers, MA). The recombinant HNRNPH1 was analyzed using an anti-FLAG antibody (F1804, Sigma-Aldrich, St. Louis, MO) and an anti-mouse HRP secondary antibody (Cell Signaling Technology, 7076).

### Chemiluminescent electrophoretic mobility shift assays (EMSA)

Gel shift assays were performed using the LightShift^TM^ chemiluminescent RNA EMSA kit (Thermo Fisher Scientific). In brief, we heated (to 95°C) and cooled 60 nM biotinylated RNA oligomer in a buffer containing 10 mM Tris-HCl (pH 7.4) as described above. In a total volume of 20 µL, 3 nM of each biotinylated RNA oligomer was incubated with recombinant HNRNPH1 protein in a binding buffer containing 10 mM Tris-HCl (pH 7.4), 1 mM EDTA, and 150 mM KCl. After 30-minute incubation at room temperature, the samples were resolved on a 6% non-denaturing polyacrylamide gel (Thermo Fisher Scientific) at 4°C, electrotransferred onto a 0.45 μm Biodyne B nylon membrane (Thermo Fisher Scientific) at 400 mA for 50 minutes at 4°C and crosslinked to the membrane (120 mJ/cm^2^, Stratalinker) for 50 seconds. The blots were then developed using the Chemiluminescent Nucleic Acid Detection Module (Thermo Fisher Scientific) and imaged on an Omega LumC (Aplegen, San Francisco, CA). To detect RNA-quadruplex shifts, we used the BG4-antibody (Absolute Antibody, Boston, MA).

### Fluorescence Resonance Energy Transfer (FRET) analysis

Fluorescence measurements were carried out using a PTI-QuantaMaster (Horiba Scientific, Piscataway, NJ) fluorometer equipped with a Peltier cell holder. The FRET-RNA oligomers in 10 mM Tris-HCl pH 7.4, 1 mM EDTA, and 25 mM KCl were denatured and reannealed by heating and cooling to room temperature as described above. Samples (120 µL) were read using a micro-fluorometer 4-side clear quartz cuvette (Starna, Atascadero, CA) with 3 mm path length. Each titration was calibrated at room temperature for 5 mins before recording the spectra. The emission spectra were obtained by setting the excitation wavelength at 500 nm and recording emission from 530 to 750 nm. Each spectrum was recorded in duplicate, averaged, and subtracted to the baseline, and plotted in Prism (GraphPad).

Spectra were normalized to the maximum emission of the donor (Cy5). The energy transfer between donor and acceptor was calculated using the parameter *P*,

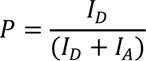

where *I_D_* is the intensity of the donor, and *I_A_* is the intensity of the acceptor (48–50). Changes in FRET efficiency were monitor as Δ*P*=*P*_final_ – *P*_initial_.

### Meta-analysis of Chromosome 22, *EWSR1*-Exon 7 – Intron 9 breakpoints reported in Ewing sarcoma primary samples and cell lines

We extracted the reported Chromosome 22 (Chr. 22) breakpoint positions or sequences from Berger *et al*., 2013 (51), Tirode *et al*., 2014 (International Cancer Genome Consortium and St. Jude’s Cloud PeCan portal (https://pecan.stjude.cloud/) (52), Brohl *et al*., 2014 (53) and Peneder *et al*., 2021 (54). Where needed, we re-mapped the reported positions from human genome Gr37 to Gr38 using the original coordinates or sequences. We focused on reported breakpoints that map to *EWSR1*-exon 7 through *EWSR1*-intron 9 and fusions involving Chromosome 11 *FLI1* translocations. For the few samples, where more than one breakpoint within *EWSR1*-exon 7 – intron 9 was reported, we used the most 5’ Chr. 22 breakpoint. The relative positions of each breakpoint within Chr. 22 were calculated and plotted in Prism (GraphPad).

### RNAi and RNA analysis

For RNAi-based experiments, cells were plated in six-well plates and transfected with 20 nM siRNA (siNeg, siHNRNPH1, or siFLI1) complexed with 4.5 µL/well Lipofectamine RNAi-Max (Invitrogen) and harvested 48 hours post-transfection. RNA isolations were performed using Maxwell 16 LEV simplyRNA purification kit (Promega). The silencing of each target transcript was confirmed using qPCR and the appropriate gene-specific primers detailed above (three independent transfections) (**Figure S1A** and **B**). To prepare libraries for sequencing, we used 300 ng of RNA with RIN > 9.5 and followed the Iso-Seq™ Express Template Preparation protocol (Pacific Biosciences, CA), selecting for transcripts above 3 kb. Full-length cDNA was synthesized and amplified using the NEBNext^®^ Single Cell/Low Input cDNA Synthesis and Amplification Module (New England Biolabs) and the Iso-Seq Express Oligo Kit (Pacific Biosciences). cDNA was amplified with a 12 cycle PCR, size-selected using 0.86 x ProNex beads (Promega) followed by an additional 3 cycle PCR. SMRTbell libraries were then prepared using the SMRTbell Express Template Prep Kit 2.0 (Pacific Biosciences). Sequencing primer v4 was annealed, and Sequel II polymerase 2.0 was bound to libraries prior to loading each on one 8M SMRT Cell on the Sequel II System using diffusion loading. Sequencing was performed with a 2-hour pre-extension and a 24-hour movie.

### Long-read sequencing

Sequence analysis of Iso-seq (PacBio) results utilized circular consensus (CCS) reads generated through sequencing. The CCS reads were run through the PacBio SMRT link version 9 IsoSeq v3 to obtain full-length, high-quality isoform consensus sequences. Briefly, for each sequencing molecule, an intra-molecular CCS read was generated. The CCS reads were then classified as full-length if a matching pair of adapter sequences was observed on the two ends of the read. The data were further refined by filtering for transcripts with poly(A) tails of at least 20 base pairs. Quality control analysis of the high fidelity reads indicated mean read lengths of about 4 kb, over 2 million reads per sample, and an estimated read quality of 99.99% accuracy. The full-length (FL) reads were run through the IsoSeq v3 pipeline that iteratively clusters FL reads belonging to the same transcript variants. Redundant sequences which have the same exon structures were collapsed into unique transcript variants with a total read count for that unique transcript variant. The unique isoform cluster reads were used to run SQANTI3 (https://github.com/ConesaLab/SQANTI3) for overall full-length transcriptome identification and quantification (55). The full-length cDNA reads were first mapped to the hg38 genome with minimap2 (56); we then used our in-house script to count the number of reads (full-length) mapped to each gene region with the coordinates from the hg38 GTF file. The raw count numbers were normalized using the trimmed mean of M values (TMM) to compare between samples. We used the longGF tool (57) to identify fusion gene reads that mapped to *EWRS1* and *FLI1* sequences. In addition, we ran the fusion_finder.py from cDNA_Cupcake package (https://github.com/Magdoll/cDNA_Cupcake) to classify and filter fusion candidates. We further manually inspected the evidence for each *EWS-FLI1* fusion transcript gene structure by mapping clustered full-length non-chimeric reads to a composite *EWSR1* and *FLI1* genomic sequence to mimic the *EWS-FLI1* fusion transcript. Results were visualized with IGV (58) and the R package Gviz (https://github.com/ivanek/Gviz). To assess the use of different 5’ transcriptional start sites or 3’ transcriptional stop sites that will define the length of the 5’ or 3’ untranslated regions (UTRs), respectively, we calculated the distance between the start of each full-length read and the end of *EWSR1*-exon 1, and the distance between the start of *EWSR1*-exon 17 and the end of each full length read. To accommodate sequencing variations, we only included clustered isoforms for which we obtained multiple reads and groups based on sizes of the respective UTR and exon of smaller or larger than 250 nucleotides (nts) to capture the differences observed in the lengths of the 5’ and 3’ UTR regions. Protein coding potential was assessed using the CPC2 web server (59). Splice strengths were calculated using the MaxEntScan algorithm using the scores generated by the Maximum Entropy Model (60) and we extracted the expression data for *EWSR1* transcript variants from the GTEx portal; https://gtexportal.org/home/gene/EWSR1.

### Minigene splicing analysis

We synthesized two DNA cassettes with 5’ *Sal1* and 3’ *Xba1* restriction site overhangs (GeneWiz, South Plainfield, New Jersey). One cassette encompassed ∼ 1000 nts of *EWSR1*-intron 7, exon 8, intron 8, and exon 9 sequences. To facilitate cloning into the pRHCGlo splicing reporter plasmid, we introduced a single nucleotide substitution into *EWSR1*-intron 8 to eliminate an *Xba1* restriction site. The second DNA cassette encompassed *EWSR1-*exon 8 and ∼1000 nts flanking sequences from *EWSR1*-introns 7 and 8 fused to ∼1700 nts from intron 5 of *FLI1* and *FLI1*-exon 6. We cloned the DNA cassettes into the pRHCGlo splicing reporter plasmid (47). We used Sanger sequencing to confirm the integrity of each plasmid clone and used the original vector pRHCGlo, pRHCGlo-*EWSR1*, and pRHCGlo-*EWS-FLI1* for subsequent cell-based studies.

We transfected HEK-293T cells (200,000 cells/well of a 6-well plate, grown overnight) with 2 µg of each plasmid, with either no other nucleic acid or 20 nM of a control siRNA (siNeg) or a siRNA targeting HNRNPH1 (siHNRNPH1) using Lipofectamine 2000 (10 µL/well plasmid DNA only; 15 µL/well plasmid DNA and siRNA). For the transfection of TC-32 cells, we used 135,000 cells/well of a 6-well plate, grown overnight, and 2 µg of each plasmid plus 20 nM siNeg or siHNRNPH1, or no additional nucleic acid complexed with Lipofectamine 3000 (10 µL/well) and the p3000 reagent (4 µL/well). Cells were grown for 48-hours, harvested, and RNA extracted (Maxwell 16 LEV simplyRNA purification kit, Promega).

First-strand cDNA synthesis was performed using 500 ng RNA, the minigene specific primer RTRHC (2 pmole), and Superscript II RT (Invitrogen) reagents at 42°C for 2 minutes. For PCR amplification, we used the 2X Phusion High-Fidelity PCR Master Mix with HF Buffer (New England Biolabs), and 300 ng of the cDNA and 10 µM of each the RSV5U and RTRHC primers, using the following thermocycling conditions - an initial denaturation at 98° C for 1 min, 27 cycles of 98° C for 10 sec, 57° C for 30 sec, and 72°C for 10 sec, followed by the final extension: 72° C for 5 min and hold at 4°C. The PCR products were separated by electrophoresis using 2% agarose gels with SYBRsafe DNA gel stain and imaged on an Omega LumC (Aplegen). Images were imported into Image J for the quantification of each amplified product. We employed the same method to study HNRNPH2 and HNRNPF using the siRNAs detailed above targeting the genes encoding each of these proteins. We confirmed the silencing efficacy of the siRNAs corresponding to *HNRNPH1*, *HNRNPH2*, or *HNRNPF* using qPCR following transfection of each plasmid/siRNA combination (three independent transfections). We normalized gene expression using the expression of *NACA* as a control. To introduce nucleotide substitutions, we replaced a ∼1.5 kb region flanked by *Nhe1* and *CsiI* restriction sites within each minigene with synthetic cassettes (GeneWiz) of the same size containing either the original sequence or one of six different sequences in which we included G-A substitutions detailed in **Figures 2D** and **S3**. The representative gel images and corresponding analysis shown are representative of at least two independent transfections.

**Figure 2:**
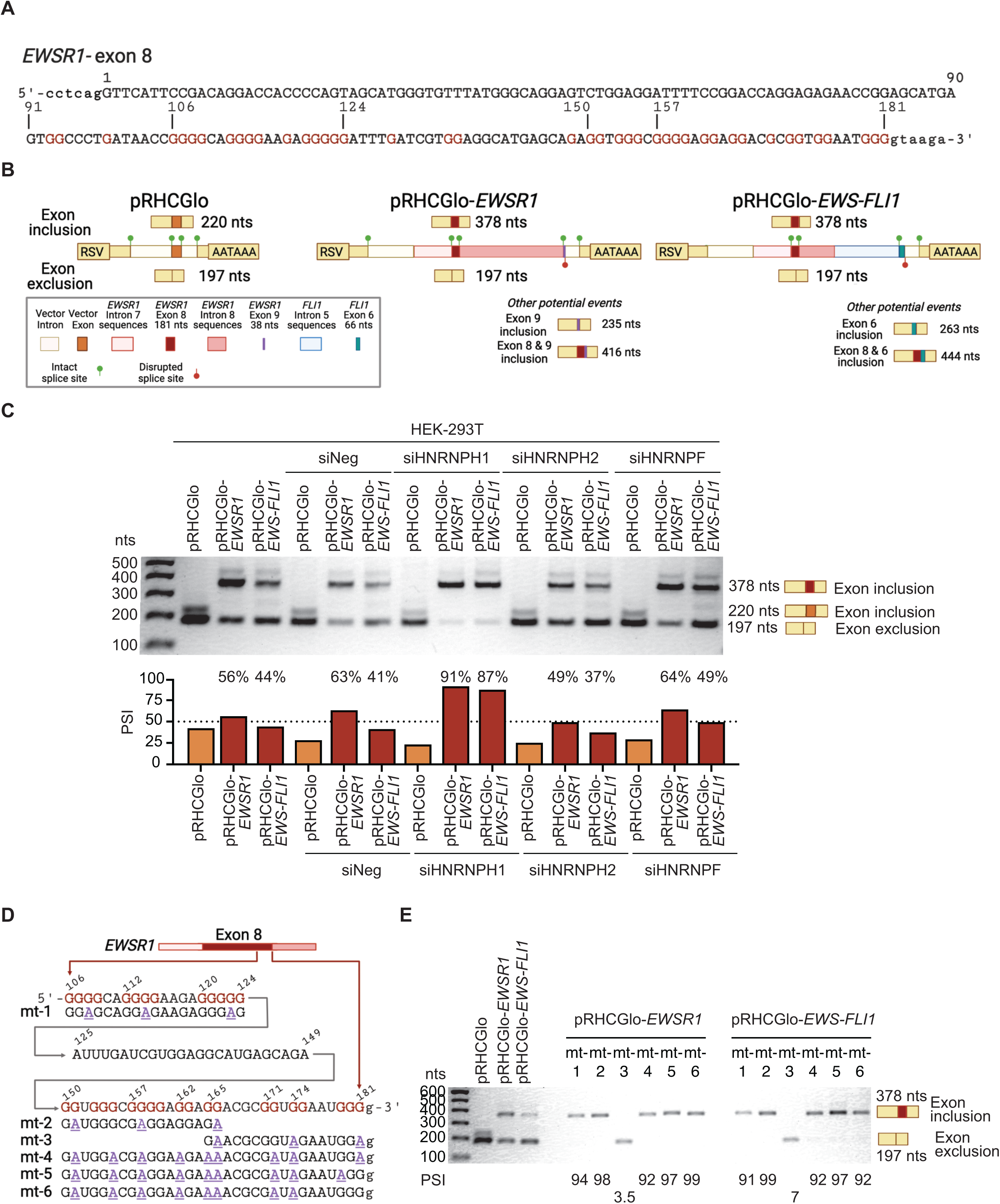
Disruption of G-tracts at the 3’ end of *EWSR1*-exon 8 alters its splicing. **A.** Schematic of *EWSR1*-exon 8 highlighting the G-rich region at the 3’ end of the exon. Lower-case letters indicate intronic and upper-case letters, exonic sequences. Numbers refer to the nucleotide coordinates of *EWSR1*-exon 8, 1 – 181 nts. **B.** Schematic of the pRHCGlo and the *EWSR1*-exon 8 centric minigenes *– EWSR1* and *EWS-FLI1 –* cloned into this vector. **C.** PCR amplified products obtained following the transfection of HEK-293T cells with the indicated plasmids or plasmids and siRNAs, and the quantification of *EWSR1*-exon 8 inclusion (percentage splice inclusion - PSI). Results are representative of at least two independent transfections of each plasmid and siRNA. **D.** Schematic of the G-A substitutions introduced into each minigene. **E.** PCR amplified products obtained following the transfection of HEK-293T cells with the indicated plasmids and the quantification of *EWSR1*-exon 8 inclusion representative of three independent transfections. **A**, **B** and **D** created in BioRender.

### G-quadruplex predictions, statistical analysis, and the visualization of protein alignments

We employed the QGRS mapper (61) to score the potential of a submitted sequence to form a G4-structure. We utilized QGRS mapper because it enables the analysis of G-run lengths of less than 3. In brief, we used the QGRS mapper to analyze the 181 nt sequence of *EWSR1*-exon 8 and 25 nt flanking intronic sequences using the following parameters, a minimum G-group length of 2, no limitation on loop size (e.g., 0 to 36), and a maximum sequence length in which a quadruplex could form of 30 nts. This analysis identified multiple overlapping sequences within the 3’ end of *EWSR1*-exon 8 predicted as capable of forming a quadruplex structure based on G-runs of 2. To prioritize guanine residues for functional analysis using minigenes or synthetic oligomers, we used the QGRS mapper to analyze the isolated G-rich sequences within the 3’ end of *EWSR1*-exon 8 and determine which nucleotide substitutions disrupt G4 formation. Statistical analysis was conducted using an unpaired t-test with Welch’s correction in GraphPad Prism 7. A p-value of <0.05 was considered significant. The protein alignments of HNRNPH1, HNRNPH2, and HNRNPF were generated using ggmsa: a visual exploration tool for multiple sequence alignment and associated data (https://github.com/YuLab-SMU/ggmsa).

## RESULTS

### The HNRNPH1-dependent processing of *EWSR1*-exon 8-*EWS-FLI1* transcripts

Ewing sarcoma cells that harbor chromosomal breakpoints within *EWSR1*-intron 8 must exclude *EWSR1*-exon 8 to express in-frame oncogenic fusion *EWS-FLI1* transcripts (**Figure 1A**). To assess the frequency of this oncogenic mutation, we conducted a meta-analysis of published studies (51–54) reporting the position of Chr22 breakpoints that map to the region encompassing of *EWSR1* exon 7 – intron 9 detected in 224 primary *EWS-FLI1*-positive Ewing sarcomas and seven Ewing sarcoma cell lines (**Figure 1B**). Our analysis determined that 36% of Ewing sarcomas harbor chromosomal breakpoints within *EWSR1*-intron 8. This percentage is higher than previously reported (62) and indicates the importance of an enhanced understanding of the splicing mechanisms used to exclude *EWSR1*-exon 8 during the maturation of the *EWS-FLI1* transcripts expressed in these tumors. Four Ewing sarcoma cell lines harbor *EWSR*1-intron 8 breakpoints, including the TC-32 cell line we selected for our subsequent cell-based analysis.

Previously, we used PCR-based analysis (9) or short-read sequencing (23) to demonstrate that the RNA binding protein HNRNPH1 is required to exclude *EWSR1*-exon 8 from *EWS-FLI1* transcripts expressed in cell lines harboring *EWSR1*-intron 8 breakpoints. However, using these methods, we could not fully determine if HNRNPH1 regulates other aspects of the processing of *EWS-FLI1* fusion transcripts or whether it functions in the biogenesis of *EWSR1* transcripts because of the challenge of distinguishing 5’ *EWSR1* and *EWS-FLI1* specific events. To overcome these limitations, we employed PacBio long-read sequencing of *EWS-FLI1* and *EWSR1* transcripts following depletion of HNRNPH1 (**Figure S1A**) or *EWS-FLI1* (**Figure S1B**) in TC-32 Ewing sarcoma cells.

**Figure 1C** shows the mapping of full-length *EWS-FLI1* transcripts identified using long-read RNA sequencing in TC-32 cells 48 hours post siRNA transfection. We also report an estimate of the relative abundance of each *EWS-FLI1* transcript variant expressed as a percentage of the total number of reads detected. **Figure 1D** shows the coverage and Sashimi plots for the mapped and clustered transcript variants of *EWS-FLI1 s*hown in **Figure 1C**, and **Table S1** details the chromosome coordinates plus other information. In control-treated cells (siNeg), 82% of full-length *EWS-FLI1* mRNAs showed evidence of *EWSR1*-exon 8 exclusion (**Figure 1C and D – left-hand panels**) with only a small proportion (4%) including this exon. In contrast, the transfection of an siRNA targeting *HNRNPH1* (**Figure 1C and D – middle panels**) resulted in a reduction in the percentage of *EWSR1*-exon 8 excluded *EWS-FLI1* mRNAs (14%) and an increase in the percentage of *EWS-FLI1* transcripts that retained *EWSR1*-exon 8 (75%). Interestingly, following the silencing of *HNRNPH1*, we also saw an overall decrease in the number of *EWS-FLI1* reads (452 versus 59) (**Figure 1C and D – middle panels**). This decrease in the number of full-length *EWS-FLI1* transcripts resembled that observed following the silencing of the fusion transcript directly); however, in this case, we detected no transcripts that retain *EWSR1*-exon 8 (**Figure 1C and D – right-hand panels**). Based on these results, we propose that the retention of *EWSR1*-exon 8 within the full-length *EWS-FLI1* transcripts reduces the overall expression of *EWS-FLI1* because the out-of-frame transcripts are the subject of nonsense-mediated decay due to the presence of the premature stop codon that generates a truncated protein (329 amino acids versus 499 amino acids: **Table S1**).

We observed no other splicing events affecting the full-length fusion oncoprotein transcripts following HNRNPH1 depletion in TC-32 cells, but we did note some alterations in transcriptional start site (TSS) usage. Specifically, in control TC-32 cells (siNeg), most *EWS-FLI1* transcripts make use of an upstream TSS, while all the *EWS-FLI1* transcripts detected in the *HNRNPH1*-silenced cells (with and without *EWSR1*-exon 8 inclusion) used a TSS closer to *EWSR1*-exon 1. We also observed that the *HNRNPH1*-silenced cells expressed a reduced variety of *EWS-FLI1* transcripts that include intronic sequences. Unlike the HNRNPH1-dependent exclusion of *EWSR1*-exon 8, neither of these changes in the complement of *EWS-FLI1* transcripts will alter the expression of the fusion oncoprotein, but we speculated that these observations could relate to the processing of *EWSR1* pre-mRNAs, so we next examined the long-read sequencing results for *EWSR1* (**Figure S1 C-E**).

Long-read sequencing of siNeg-transfected TC-32 cells (**Figure S1C**) identified two dominant full-length *EWSR1* transcripts that align with the consensus sequences of *EWSR1-206* (ENST00000397938) (21% of reads) and *EWSR1-207* (ENST00000406548) (30% of reads). Both *EWSR1-206* and *EWSR1-207* consist of 17 exons but utilize slightly different transcriptional start and stop sites and contain 38 or 35 nucleotide versions of exon 9, respectively. Interestingly, we observed no disruption in *EWSR1* exon 8’s inclusion in the absence of HNRNPH1, but we did detect fewer transcript variants expressing the 35-nucleotide form of exon 9, with most now including the 38-nucleotide form of this exon (**Figure S1D and Table S2**). Based on these findings, we speculate that in the context of the full-length *EWSR1* transcript, *cis*-acting sequences counter the HNRNPH1-dependent exon 8 exclusion event observed in the context of the *EWS-FLI1* transcript, but it may function in mediating the splicing event that generates the 35 nts exon 9 expressed by the *EWSR1-207* transcript variant. Critically, these observations suggest that targeting the HNRNPH1-dependent processing of *EWS-FLI1* should still maintain EWSR1 protein expression, as the isoforms expressed by EWSR1-206 (isoform 1 - Q01844-1 656 amino acids) and EWSR1 207 (isoform 3 - Q01844-3 655 amino acids) differ by only one amino acid at position 326. We also saw no evidence for HNRNPH1’s function in expressing two rare *EWSR1* variants *(EWSR1-201,* ENST00000331029*, EWSR1-203,* ENST0000033205) that both involve alternative splicing events focused on *EWSR1*-exon 8 that we previously discussed as possibly involving HNRNPH1 function for their expression (23). However, long-read sequencing did detect other transcript variants in both siNeg and siHNRNPH1 samples, albeit making up less than 1% of all reads. These transcript variants include those exhibiting exon 6 or exon 9 exclusion events and variations in the sequences forming exon 15, all of which maintain protein-coding potential (**Table S2**).

Consistent with our analysis of *EWS-FLI1* transcripts, we observed that the silencing of *HNRNPH1* resulted in the use of different regulatory signals up and downstream of the consensus *EWSR1* coding sequences (**Figure S1F**). Furthermore, we observed a substantial shift in the proportion of *EWSR1* transcripts containing intronic sequences following the silencing of *HNRNPH1* (48% in control and 67% in siHNRNPH1-transfected cells) but with reduced sequence diversity of intron-containing transcripts compared to control cells (**Figures S1C and S1D)**. Interestingly, most of the full-length *EWSR1* transcripts that are non-protein-coding contain intron 7 and/or intron 8 sequences under all conditions. It is unclear why the *EWSR1* gene expresses such a substantial proportion of non-protein-coding transcripts versus protein-coding transcripts, but our data suggest that HNRNPH1 may function in regulating the overall abundance and diversity of these sequences. Overall, we observed fewer differences in the abundance and make-up of *EWSR1* transcripts following the silencing of *EWS-FLI1* compared to control transfected cells, but these changes included the detection of full-length *EWSR1* transcripts that included a longer 5’-UTR and a shorter 3’-UTR than those previously annotated (**Figures S1E and S1F**).

Using the meta-analysis of *EWSR1* breakpoints and our long-read sequencing of the *EWS-FLI1* and *EWSR1* transcripts as guides, we next generated minigene constructs to analyze potential *cis*-acting sequences within *EWSR1*-exon 8 that contribute to determining its inclusion or exclusion within a particular sequence context.

### Mutation of G-rich sequences within the 3’ end of *EWSR1*-exon 8 alters the inclusion of this exon

*EWSR1*-exon 8 is 181 nts in length (**Figure 2A**). The predicted Maximum Entropy Model scores for the *EWSR1*-Exon 8/Intron-8 5’ splice site is 7.83, a score that falls within the intermediate category of 5’ splice site strength, suggesting that adjacent sequences can influence splice site selection. Intronic G-rich sequences can act as Intron Splice Enhancers (ISEs), while G-tracts within exons can act as Exon Splice Silencers (ESSs), inducing exon skipping or altering splice site selection (63). The first GGG within *EWSR1*-intron 8 is 66 to 69 nts from the last nucleotide of *EWSR1* exon 8 (**Figure S2A**), consistent with previous studies highlighting enrichment of such G-tracts within 70 nts of low or intermediate strength 5’ splice sites (10). We developed two minigene systems to investigate the HNRNPH1-mediated processing of transcripts containing *EWSR1*-exon 8 (**Figure 2B**).

In brief, we cloned two DNA cassettes into the pRHCGlo splicing reporter plasmid (47). One DNA cassette consisted of the *EWSR1*-intron 7, exon 8, intron 8, and exon 9 sequences. The second DNA cassette consisted of *EWSR1-*exon 8 and ∼1000 nts flanking sequences from *EWSR1*-introns 7 and 8 fused to ∼1700 nts from intron 5 of *FLI1* and *FLI1*-exon 6. This design considered the need to encompass the region that harbors most of the documented breakpoints in *EWSR1*-intron 8 observed in Ewing sarcomas (**Figure 1B**) and a fused intron of a size comparable to that of the wild-type *EWSR1*-intron 8.

We retained the endogenous splice sites flanking *EWSR1-*exon 8 in both constructs and the 3’ splice acceptor sites adjacent to *EWSR1*-exon 9 or *FLI1*-exon 6 in the respective constructs, but not the 5’ splice donor sites. To assess exon inclusion versus exclusion events, we used a plasmid-specific primer (RTRHC) to initiate reverse transcription and amplified the cDNA using a 5’ PCR primer corresponding to sequences within the RSV region of the plasmid (RSV5U) and the RTRHC oligomer as the 3’ PCR primer. **Figure 2B** shows a schematic of the predicted amplified products that these primers should detect.

We first transfected the minigene constructs (pRHCGlo-*EWSR1* and pRHCGlo-*EWS-FLI1*) into HEK- 293T cells alone or combined with a negative control siRNA (siNeg) or siRNAs targeting *HNRNPH1* (siHNRNPH1), *HNRNPH2* (siHNRNPH2), or *HNRNPF* (siHNRNPF), and then harvested RNA 48 hours later (**Figure S2B**). The gels and accompanying quantification indicate that HNRNPH1 depletion results in a substantial increase in *EWSR1*-exon 8 inclusion in the context of the pRHCGlo-*EWSR1* and pRHCGlo-*EWS-FLI1* constructs (**Figure 2C**). Consistent with our previous findings (23), depletion of HNRNPH2 or HNRNPF did not alter the splicing of *EWSR1*-exon 8, despite their homology to HNRNPH1, indicating the specificity of HNRNPH1’s function in the regulation of certain splicing events.

Under control conditions (no siRNA or siNeg co-transfection), we observed slight differences in the percentage inclusion of *EWSR1*-exon 8 when present in the context of adjacent *EWSR1* intronic sequences (∼60%) versus the artificial EWS-FLI1 fusion intron (∼40%). However, silencing of *HNRNPH1* resulted in similar percentages of *EWSR1*-exon 8 inclusion irrespective of the adjacent sequences. Based on these findings, we speculate that HNRNPH1 is necessary to exclude *EWSR1*-exon 8, but *cis*-acting sequences that differ between the two constructs contribute to the inclusion of the exon. Interestingly, the differences between the two minigene constructs were less pronounced when we conducted the same studies in the TC-32 Ewing sarcoma cell line transfections (**Figure S2C**). However, this may reflect the reduced transfection efficiency of the TC-32 cells versus HEK-293T cells, and as expected, we still observed the increase in *EWSR1*-exon 8 inclusion in cells in which we also depleted HNRNPH1.

We have demonstrated using RNA oligomers that HNRNPH1 can bind the G-rich sequences towards the 3’ end of *EWSR1*-exon 8 (**Figure 2A**) and that at least one of these sequences (nts 103 – 124 of the exon) can form a G4 structure (23). To examine the contribution of the G-rich sequences within *EWSR1-* exon 8’s 3’ end to the inclusion or exclusion of this exon in cell-based studies, we modified our minigene constructs to incorporate nucleotide substitutions designed to disrupt G-tracts that HNRNPH1 could bind or that could contribute to RNA folding through non-Watson-Crick base pairing. In brief, we replaced a ∼1.5 kb region within each minigene with synthetic cassettes of the same size containing either the original sequence or one of six mutant sequences (mt-1 – mt-6) in which we included G-A substitutions which could disrupt HNRNPH1 binding and/or RNA structure predicted using the QGRS mapper algorithm (47) (**Figure 2D** and **Figure S3**). **Figure 2E** shows a representative image of the inclusion or exclusion of *EWSR1*-exon 8 using the control and mutant minigenes.

The first G-A substitutions (mt-1) altered the sequence of the region we studied previously (nts 106 – 124) (23). Consistent with our earlier findings, disruption of the three G-tracts within this sequence results in *EWSR1*-exon 8 inclusion (>90%) and highlights this region as a lead candidate *cis*-regulatory element defining the splicing of *EWSR1*-exon 8 (23). Next, we considered a 32 nt region (150 – 181 nts) at the 3’ end of *EWSR1*-exon 8. We have previously shown that HNRNPH1 can bind RNA oligomers that include the longest G-tracts within this region (nts 153 – 173). However, the density of the G-tracts within this region is such that it is challenging to prioritize residues for analysis in cell-based systems, as there are several sequences that HNRNPH1 could bind and many combinations of G-residues that could contribute to a predicted G4 structure (**Figure S3**). Nevertheless, we first examined the effect of G-A substitutions predicted to disrupt G4 structures formed by nucleotides 150 through 166 (mt-2) and nucleotides 166 through 181 (mt-3). These sequence substitutions generated dramatically different results, with the first showing ∼98% inclusion of *EWSR1*-exon 8, and the latter, less than 10%, confirming that within this region are sequences that could function as a *cis*-regulatory element defining inclusion or exclusion of *EWSR1*-exon 8. Next, we generated minigenes in which we made G-A substitutions predicted to disrupt the formation of G4 structures that considers the entire 150 – 181 nt region (mt-4, mt-5, and mt-6). These minigenes all showed *EWSR1*-exon 8 inclusion (>90%). As part of the design of these more extensively modified minigenes, we considered if the G-A substitution at the end of mt-3 represented a critical residue determining exon skipping, irrespective of the introduction of other substitutions. Interestingly, we observed that a G-A substitution of nt 181 (or nt 179), did not counter the effect of the other substitutions resulting in *EWSR1*-exon 8 inclusion. In summary, our results indicate multiple G-residues throughout the 150 - 181 nt of *EWSR1*-exon 8 have the potential to contribute to the regulation of this exon’s splicing, and based on these results, we next selected to examine the *in vitro* biophysical properties of RNA oligomers that included the G-tracts present between 100 and 124 nts of *EWSR1*-exon 8, and two overlapping RNA oligomers encompassing the region 150 – 181 nts.

### Multiple G-rich sequences at the 3’ end of *EWSR1*-exon 8 form G4 structures

To study the RNA structure of the *EWSR1*-exon 8 G-tracts that included those which when mutated, altered the splicing of this exon, we synthesized a series of RNA oligomers we refer to as rG1 (nts 103 – 127 of *EWSR1*-exon 8), rG2 (nts 150-178), and rG3 (nts 152 - 181) (**Figure 3A**) based on the results of the minigene studies and our previous analysis of the rG1 and rG2 sequences (23). G-rich sequences can, in the presence of a monovalent cation, form G4 structures, and **Figure S3** summarizes the G4 prediction values (61) of these sequences. We assessed the *EWSR1*-exon 8 RNA oligomers’ structure by examining their CD spectra under biologically relevant salt concentrations (150 mM KCl). The three RNA oligomers exhibited similar CD spectra - a negative peak intensity at 240-245 nm and a positive peak intensity of 262-265 nm, indicating these sequences form parallel (5’-3’) G4 structures (**Figure 3B**) (64, 65). The spectrum for rG1 is consistent with our previous analysis (23). Furthermore, the rG2 and rG3 results align with the transcriptome-wide analysis of rG4s that reported reverse transcriptase stalling at the 3’ end of *EWSR1*-exon 8 (15, 16). Supporting our CD spectra results, computational analysis of the rG1, rG2, and rG3 sequences (61) indicates the potential for the RNA oligomers corresponding to these G-rich regions forming G4 structures, but only if using G-runs of 2 nucleotides **(Figure S3)**, which precludes the formation of three-tier G4s, and instead suggests the formation of two-tier quartet structures such as modeled in **Figure 3C**. Of note, we previously observed enrichment of G-rich motifs predicted to form two-tier quartet G4s in exonic sequences bound by HNRNPH1 (23). In the case of the rG1, rG2, and rG3 sequences, depending on the guanine residues that form the G-quartet, these two-tier G4-structures could exhibit polymorphism with variations in the length and nucleotide composition of the loops connecting the G-tracts (**Figure S3**).

**Figure 3:**
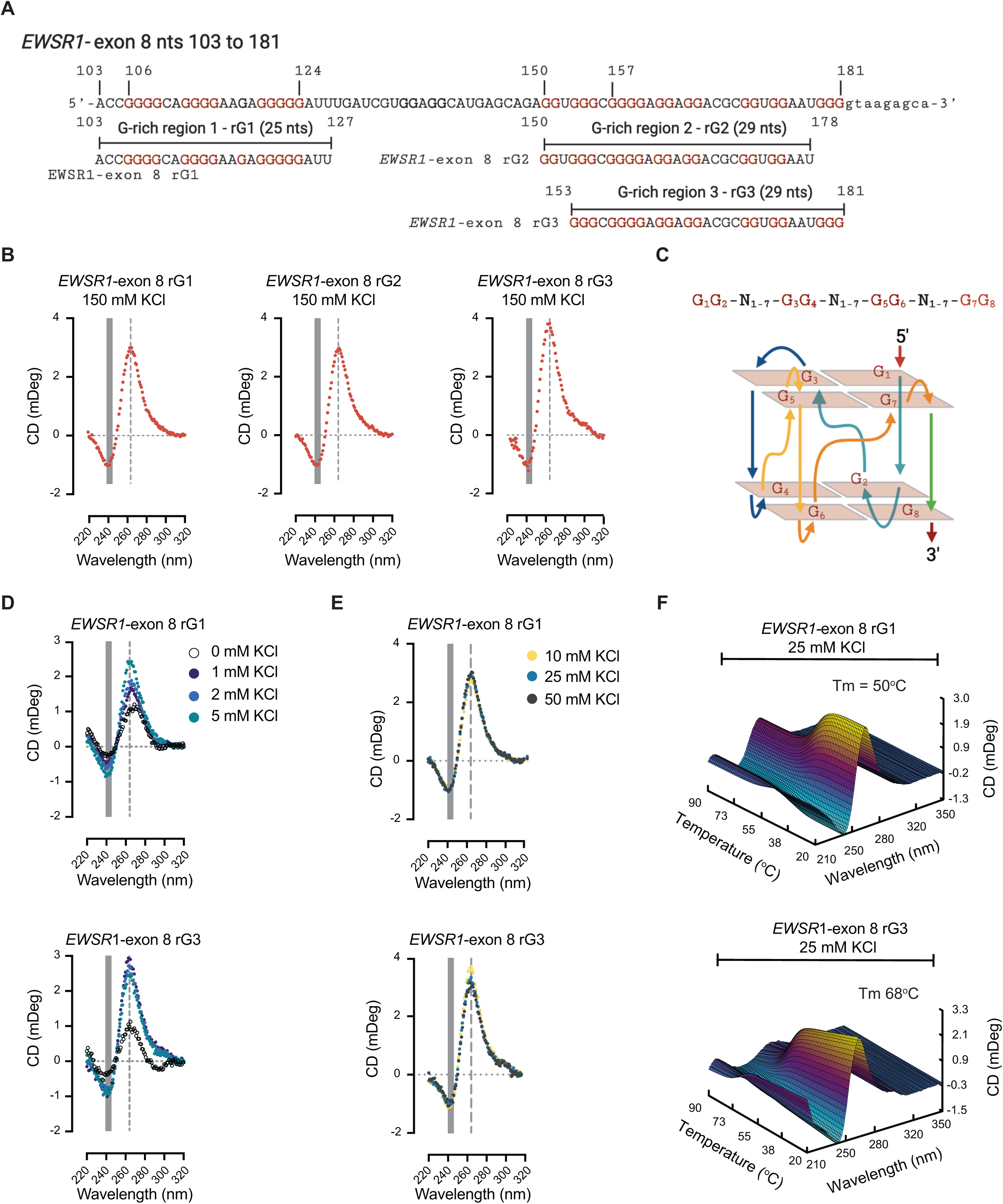
Multiple G-rich sequences at the 3’ end of *EWSR1*-exon 8 form G4 structures. **A.** Schematic of the 3’ end of *EWSR1*-exon 8 highlighting the G-rich regions and the sequences of RNA oligomers used in this study. Red Gs indicate those guanines within G-tracts of two or more nucleotides. **B.** CD spectra for the indicated RNA oligomers (150 mM KCl). **C.** A schematic model of a predicated two-tier quartet parallel G4-structure based on the rG1, rG2, or rG3 sequences formed in the presence of a monovalent cation (not shown). **D.** CD spectra for the indicated RNA oligomers in the presence of no added KCl, 1, 2, or 5 mM KCl. **E.** CD spectra for the indicated RNA oligomers in the presence of 10, 25, or 50 mM KCl. **F.** Thermal melt curves for the indicated RNA oligomers (25 mM KCl). **A** and **C** created in BioRender.

To evaluate the *EWSR1*-exon 8 RNA oligomers’ propensity to form G4-structures and stability once folded, we determined the minimum salt concentration that generated CD spectra consistent with G4- formation and their thermal melting (T_m_) points. We observed shifts in the CD spectra at 240-245 nm with the most substantial shift at 260-265 nm, compatible with G4 formation for each oligomer at salt concentrations as low as 1 – 5 mM KCl (**Figure 3D** and **Figure S4A**). At concentrations between 10 – 50 mM KCl, the CD spectra (**Figure 3E** and **Figure S4B**) resembled those seen at 150 mM KCl (**Figure 3B**). T_m_ curves generated for the rG1 oligomer demonstrated a T_m_ value of 50°C at 25 mM KCl (**Figure 3F**). At 25 mM KCl, the rG2 (**Figure S4C**) and rG3 RNA oligomers (**Figure 3F**) exhibited even higher melting points, which, in the case of rG3, exceeded 60°C. Interestingly, the rG1 oligomer exhibited a gradual change in the maxima at 260-265 nm between 1 and 5 mM KCl, whereas at KCl concentrations as low 1 mM, the CD spectra of the rG3 oligomer resembled that observed at 150 mM KCl. These observations suggest the rG1 oligomer may form intermediate states, but critically, in the presence of physiologically relevant concentrations of monovalent ions, the energy barrier to forming rG4 structures is low for all three RNA oligomers (rG1, rG2, and rG3).

Of note, the higher T_m_ values observed for rG3 indicate that rG1 and rG3 quadruplex structures may differ in loop compositions despite having similar parallel structures. Analysis of the thermal melting of rG1, rG2, or rG3 at RNA concentrations of 10 - 30 µM generated similar melting values (**Figures S4D** – **F**), indicating that in the case of these RNA oligomers, quadruplex formation is independent of RNA concentration. Collectively, these results suggest that any transcripts containing these endogenous *EWSR1* sequences are likely to form G4 structures under physiological relevant cationic salt conditions rather than remain in a non-G4 state unless an active mechanism exists to inhibit the generation of such structures or alter them once formed. Therefore, we hypothesized that in cells, altering the secondary folding of the nascent RNA at the 3’ end of *EWSR1*-exon 8 will require interaction with an RNA-binding protein or proteins (RBPs) such as HNRNPH1.

### RNA structure determines the kinetics of HNRNPH1 binding to *EWSR1*-exon 8 G rich sequences

To examine HNRNPH1’s binding to *EWSR1*-exon 8, we focused on the rG1 and rG3 sequences. We selected to study these sequences because though they both form G4 structures, they exhibit substantial differences in their T_m_ (50°C versus 68°C), enabling us to assess if this alters the consequence of HNRNPH1 binding. For these studies, we examined wild-type versions of the rG1 and rG3 sequences and two different versions of each sequence designed to disrupt the RNA secondary structure (**Figures 4A and B**). In one set, we made G-A substitutions (rG1-mt and rG3-mt) to disrupt the G-tracts, while in the second set (rG1-7d-G and rG3-7d-G), we substituted some of the guanosines with 8-aza-7- deazaguanosines (Super-G, IDT). 8-aza-7-deazaguanosines substitutions retain the original oligomer’s sequence but eliminate the Hoosgteen hydrogen bonds needed for G-quadruplex formation. We selected which residues to alter based on the predicted G4 structures for both sequences (**Figures S3A** and **B**) and the results of the mutant minigene studies. CD spectra of the RNA oligomers in which we made G-A substitutions (rG1-mt and rG3-mt) indicated these sequences form an A-form-like structure with minima at 250 nm and maxima at 270 nm and no change in spectra in the presence of 150 mM KCl (66). The CD spectra of the 7d-G modified oligomers also confirmed disruption of the generation of G4 structures (**Figure 4B**). Specifically, their spectra closely resemble those of the unmodified sequences at 0 mM KCl, and in agreement with previous studies, the addition of KCl resulted in no evidence of spectra consistent with the formation of a G4 structure (67, 68).

**Figure 4:**
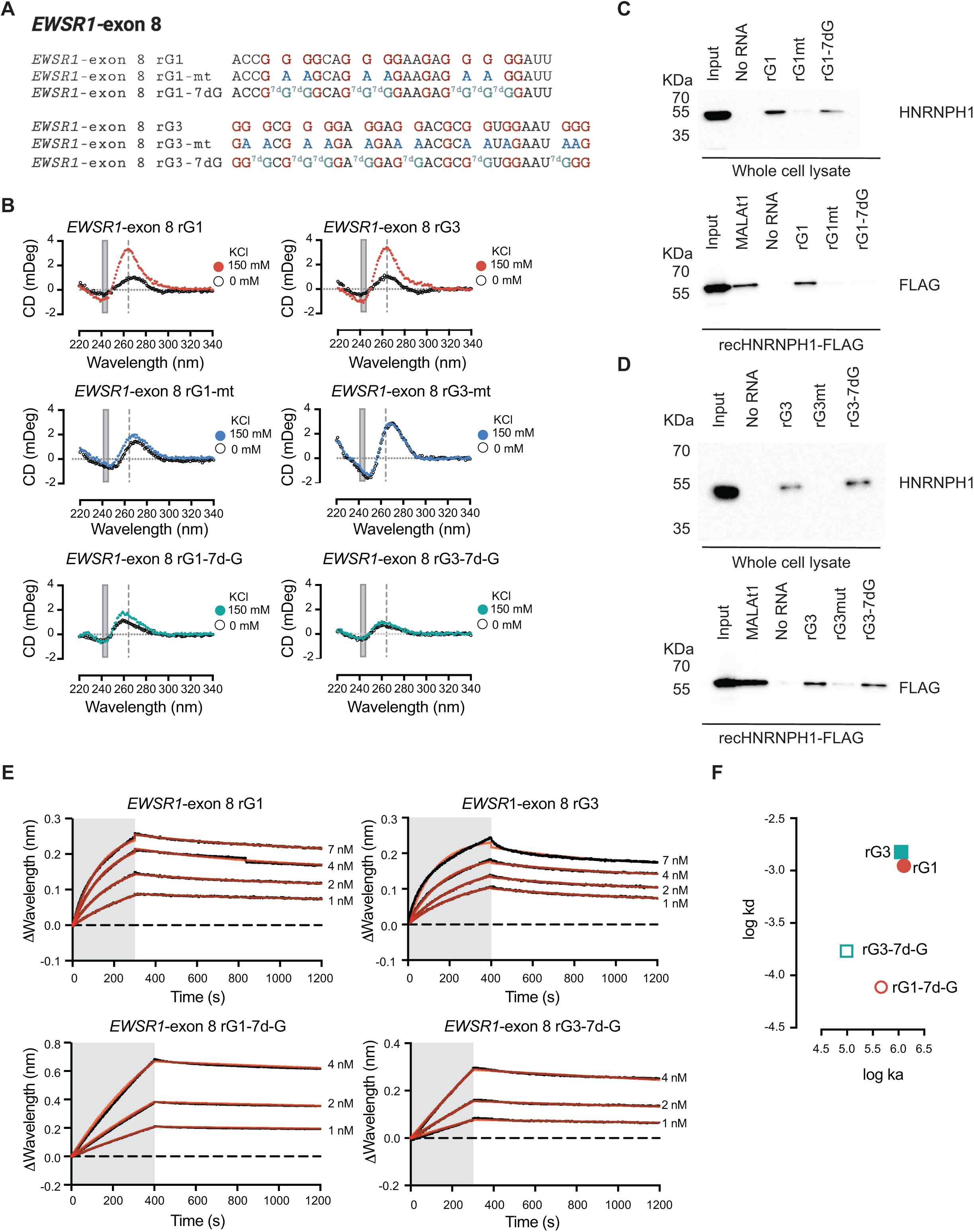
RNA structure determines the kinetics of HNRNPH1 binding to *EWSR1*-exon 8 G rich sequences. **A.** The sequences of the rG1 and rG3 RNA oligomers and the G-A and 8-aza-7-deazaguanosines (7d-G) variants. **B.** CD spectra for the indicated RNA oligomers in the presence of no added KCl or 150 mM KCl. **C.** RNA-pulldowns for rG1 using whole-cell lysate prepared from TC-32 cells or FLAG-tagged recombinant HNRNPH1. **D.** RNA-pulldowns for rG3 using whole-cell lysate prepared from TC-32 cells or FLAG-tagged recombinant HNRNPH1. **E.** Sensorgrams for the indicated RNA oligomers and HNRNPH1 at increasing protein concentrations. **F.** Graphical representations of the association and disassociation values of the indicated RNA oligomers. **C** and **D.** The RNA-pulldowns and immunoblots are representative of at least two independent experiments. **A** created in BioRender.

We first employed RNA pull-downs using the biotin added to each oligomer at the 5’ end and whole-cell lysate from TC-32 Ewing sarcoma cells or recombinant FLAG-tagged HNRNPH1 to assess the binding of HNRNPH1 to rG1 and rG3. Before adding RNA to whole-cell lysate or recombinant protein, we incubated each oligomer at 95°C and cooled the RNA to room temperature in the presence of 150 mM KCl (**Figures 4C and D**). Immunoblotting for HNRNPH1 demonstrated binding to the rG1 and rG3 oligomers, but consistent with HNRNPH1’s recognition of specific sequences, we observed much-reduced binding of the rG1-mt or rG3-mt oligomers. The results for the binding of the rG1-7d-G versions of these sequences using whole-cell lysates indicated binding of the rG1-7d-G and rG3-7d-G oligomers by HNRNPH1, but we only observed reproducible binding of the rG3-7d-G using the HNRNPH1 recombinant protein. To assess if this reflected a measurable difference in binding or a lack of sensitivity of the antibody-based assay, we next utilized a more sensitive and quantitative approach, Biosensor Bio-layer Interferometry (BLI), to investigate the binding of HNRNPH1 to the rG1 and rG3 sequence and the rG1-7d-G and rG3-7d-G variants (**Figure 4E**).

BLI analysis makes use of biotinylated RNA oligomers tethered to a 3D streptavidin surface over which we flowed increasing concentrations of recombinant HNRNPH1. The biosensor detects the interaction between protein and RNA as a change in the interference pattern of light reflected from the sensor surface and an internal reference layer, expressed as wavelength. Our real-time analysis allowed us to measure kinetic parameters that define the binding affinity of HNRNPH1 for each RNA oligomer (**Table 1**). HNRNPH1’s binding of rG1 determined an equilibrium dissociation constant K_D_ of 1.08 ± 0.53 nM; findings of a similar magnitude as determined previously by biosensor surface plasmon resonance (SPR) analysis (23). Analysis of HNRNPH1’s binding of rG3 measured a K_D_ of 1.39 ± 0.54 nM. HNRNPH1 also bound the rG1-7dG and rG3-7dG RNA oligomers with sub-nanomolar affinities, suggesting that the differences observed using the RNA-pulldown method reflect the lack of sensitivity of the antibody-based assay. Additional BLI analysis determined that HNRNPH1 also binds a folded RNA oligomer corresponding to the rG2 that overlaps with the rG3 sequence with a low nanomolar affinity (K_D_ of 3.1± 0.7 nM (**Figure S4G**), while rG1-mt and rG3-mt showed weak binding interactions between HNRNPH1 and RNA (**Figure S4H**), agreeing with the antibody-based assay (**Figure 4C** and **D**).

**Table 1.** BLI kinetic and thermodynamic parameters for the interaction of HNRNPH1 with *EWSR1*-exon 8 rG1 and rG3 RNA oligomers

|  | **K<sub>D</sub> (M)** | **K<sub>D</sub> error** | **k<sub>a</sub> (1/Ms)** | **k<sub>a</sub> error** | **k<sub>d</sub> (1/s)** | **k<sub>d</sub> error** |
| --- | --- | --- | --- | --- | --- | --- |
| **rG1** | 1.08E-09 | 5.33E-10 | 1.27E+06 | 6.34E+05 | 1.11E-03 | 2.43E-04 |
| **rG1-7d-G** | 2.06E-10 | 7.37E-11 | 4.61E+05 | 5.30E+03 | 7.73E-05 | 6.93E-07 |
| **rG3** | 1.39E-09 | 5.43E-10 | 1.13E+06 | 8.24E+05 | 1.51E-03 | 2.98E-04 |
| **rG3-7d-G** | 1.78E-09 | 5.06E-11 | 9.79E+04 | 2.78E+03 | 1.70E-04 | 9.20E-07 |

Interestingly, though the BLI-determined low nM binding affinities proved similar for the rG1, rG3, and 7dG variants, the kinetic binding parameters showed that the association and dissociation rates for the interactions between HNRNPH1 and the G4 and non-G4 (7dG) RNAs differ, with both rates faster for RNA in a G4-state than when the same sequence is in a non-G4 state (**Figure 4F, Table 1**). Specifically, the association rate kinetic of the protein is faster by 3-fold to rG1 and 11-fold to rG3, while the dissociation rate kinetic is faster by more than 10-fold for both the non-modified sequences compared to the 7dG RNA oligomers. An interpretation of these results is that HNRNPH1 recognizes G-rich sequences in a G4-state faster than those in a non-G4 conformation. However, the population of HNRNPH1 bound to the latter state will dominate over time since the dissociation rate of protein to the non-G4 state is slower than that observed for the same RNA in a G4 state. HNRNPH1’s faster recognition of G4 structured RNAs may also mitigate further RNA folding.

### Mobility shift analysis confirms HNRNPH1’s binding of RNA oligomers corresponding to *EWSR1*- exon 8 G-rich sequences when in a G4 or non-G4 state

To confirm and extend our study of HNRNPH1’s binding of RNA oligomers corresponding to *EWSR1*- exon 8 G-rich regions, we next examined native-gel electrophoretic mobility shifts of the rG1, rG3, and their respective 7dG variants (3 nM each) using a G4-specific antibody (BG4) in the presence of 150 mM KCl (**Figure 5A**). Under these conditions, the rG1 and rG3 oligomers exhibited electrophoretic mobility shifts when complexed with the BG4 antibody, consistent with the formation of RNA-protein complexes with minimal mobility.

**Figure 5:**
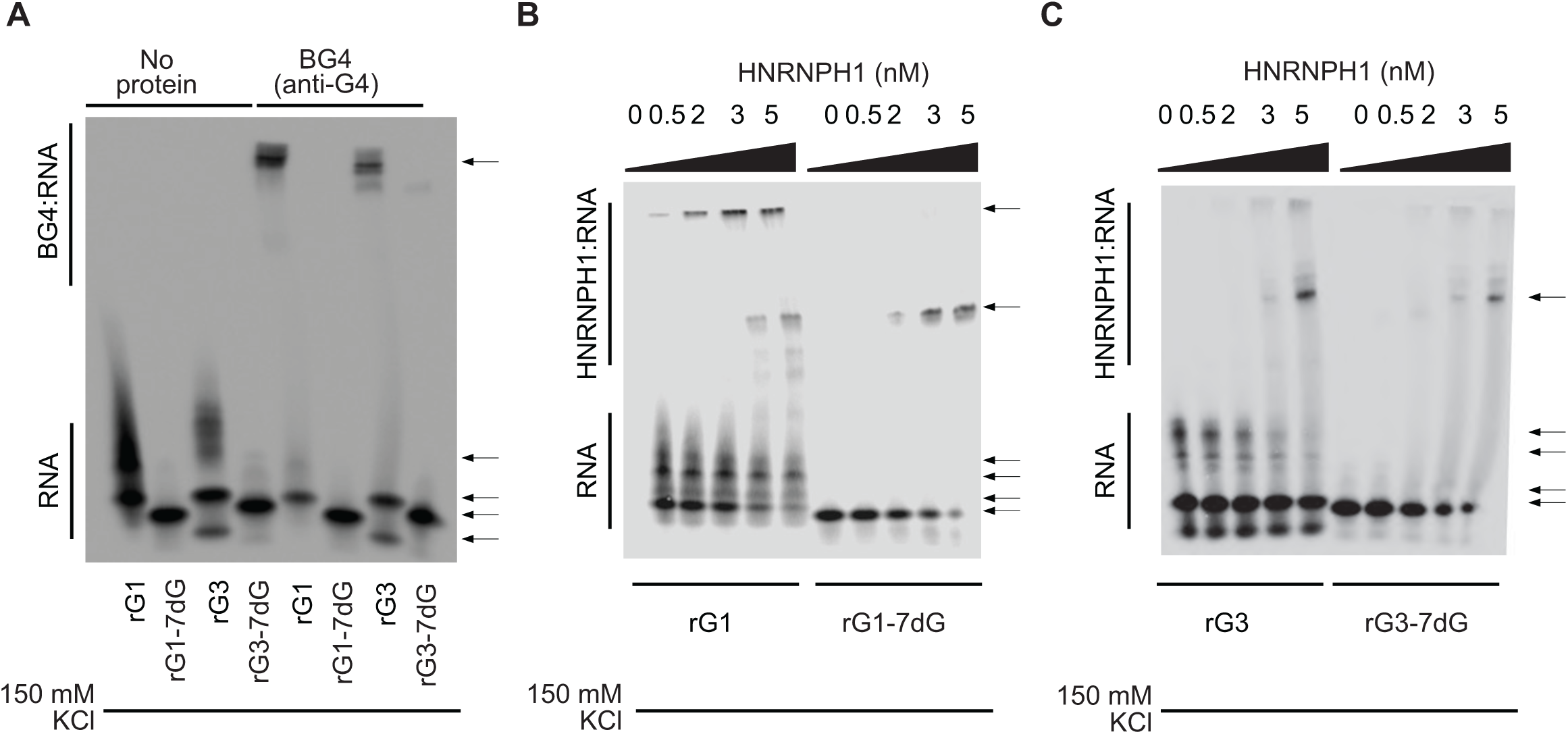
Mobility shift analysis confirms HNRNPH1’s binding of RNA oligomers corresponding to *EWSR1*-exon 8 G-rich sequences when in a G4 or non-G4 state. **A.** EMSA-based analysis of the rG1 and rG3 RNA oligomers and their respective 7d-G variants in the absence or presence of the BG4 antibody. **B.** EMSA-based analysis of the rG1 and rG1-7d-G RNA oligomers in the absence or presence of increasing concentrations of HNRNPH1. **C.** EMSA-based analysis of the rG3 and rG3-7d-G RNA oligomers in the absence or presence of increasing concentrations of HNRNPH1. Each EMSA gel shown is representative of at least two independent experiments.

We next examined the electrophoretic mobility shifts rG1 or rG3 and their respective 7dG mutants incubated with increasing amounts of HNRNPH1 (0 – 5 nM) (**Figures 5B and C**). When run through a gel matrix, the population of unbound rG1 RNA runs as multiple species indicative of the formation of higher-order structures. Such higher order structures can result from the stacking of intramolecular or intermolecular G4s, with the CD thermal melting studies (**Figures S4D and S4F**) indicating intramolecular formation.

At low HNRNPH1 (0.5 and 2 nM) concentrations, we detected HNRNPH1-rG1 gel shifts, consistent with HNRNPH1’s interaction with RNA in a form that resulted in a greater shift than that seen when bound to the rG1-7dG oligomer (**Figure 5B**). Interestingly, at higher concentrations (3 and 5 nM), we observed the same large shift and a shift that ran at the same mobility of the complexes formed when we analyzed the same concentration of HNRNPH1 bound to the rG1-7dG RNA oligomer. We detected no rG3 gel shifts at low HNRNPH1 concentrations, but at 3 and 5 nM HNRNPH1, we observed two gel shifts, one that, as with rG1, exhibited little mobility and a second that exhibited a mobility shift comparable to that of the HNRNPH1:rG3-7dG complexes (**Figure 5C**), though in both cases, the signal was less intense than that detected for the rG1 oligomer. The detection at the two higher protein concentrations of HNRNPH1-rG1 or HNRNPH1-rG3 complexes exhibiting the same mobility as observed for the RNA-protein complexes formed with their respective 7dG oligomers could reflect binding at the higher protein concentrations of any remaining RNA in a non-G4 state within each sample. An alternative hypothesis is that at higher HNRNPH1 concentrations, a proportion of the HNRNPH1-RNA complexes formed results in a structural change that generates a complex with mobility akin to that observed for the complexes formed when HNRNPH1 binds the 7dG oligomer. We thus next used spectroscopic methods to investigate this latter possibility.

### HNRNPH1 destabilizes the G4 structures formed by *EWSR1*-exon 8 rG1 and rG3 RNA oligomers

We synthesized double-terminal fluorescence-labeled oligomers modeled after rG1 and rG3 (rG1-FRET and rG3-FRET) labeled at the 5’ end with Cy5 (acceptor) and Cy3 (donor) at the 3’ end (**Figure 6A**) and confirmed that these modified RNAs form G4 structures at 150 mM KCl (**Figure 6B**). Bulk salt titration shows the normalized intensity of the acceptor, Cy5, at 662 nm increases proportionally to the concentration of KCl (1 - 150 mM KCl) (**Figure 6C** and **Figure S5A**). Calculating the energy transfer between the donor and acceptor (*P)* as described in the *Material and Methods*, we determined that the rG1-FRET and rG3-FRET RNA oligomers at 150 mM KCl (G4 state) have *P* values of 0.40 and 0.38, respectively. In comparison, in the presence of no added salt (non-G4-state), we determined *P* values for the rG1-FRET and rG3-FRET RNA oligomers as 0.50 and 0.45, respectively. Comparing these values indicated that altering the salt concentration generates a Δ*P* of 0.1 for rG1-FRET and 0.07 for rG3 FRET (0 versus 150 mM KCl). These energy transfers from the acceptor to the donor indicate that the pair of fluorophores are closer in space at increasing concentrations of KCl, indicative of a conformational change from a non-G4 state to a compact G4 structure (52, 54). Consistent with the CD results (**Figure 3D**), the rG1 and rG3-FRET signals saturate at KCl concentrations of 10 mM or higher (**Figure S5A**). We propose that the slight differences between the behavior of the rG1 and rG3 FRET oligomers relate to the differences in the size of the two RNAs (rG1 - 25nts; rG3 – 29 nts).

**Figure 6:**
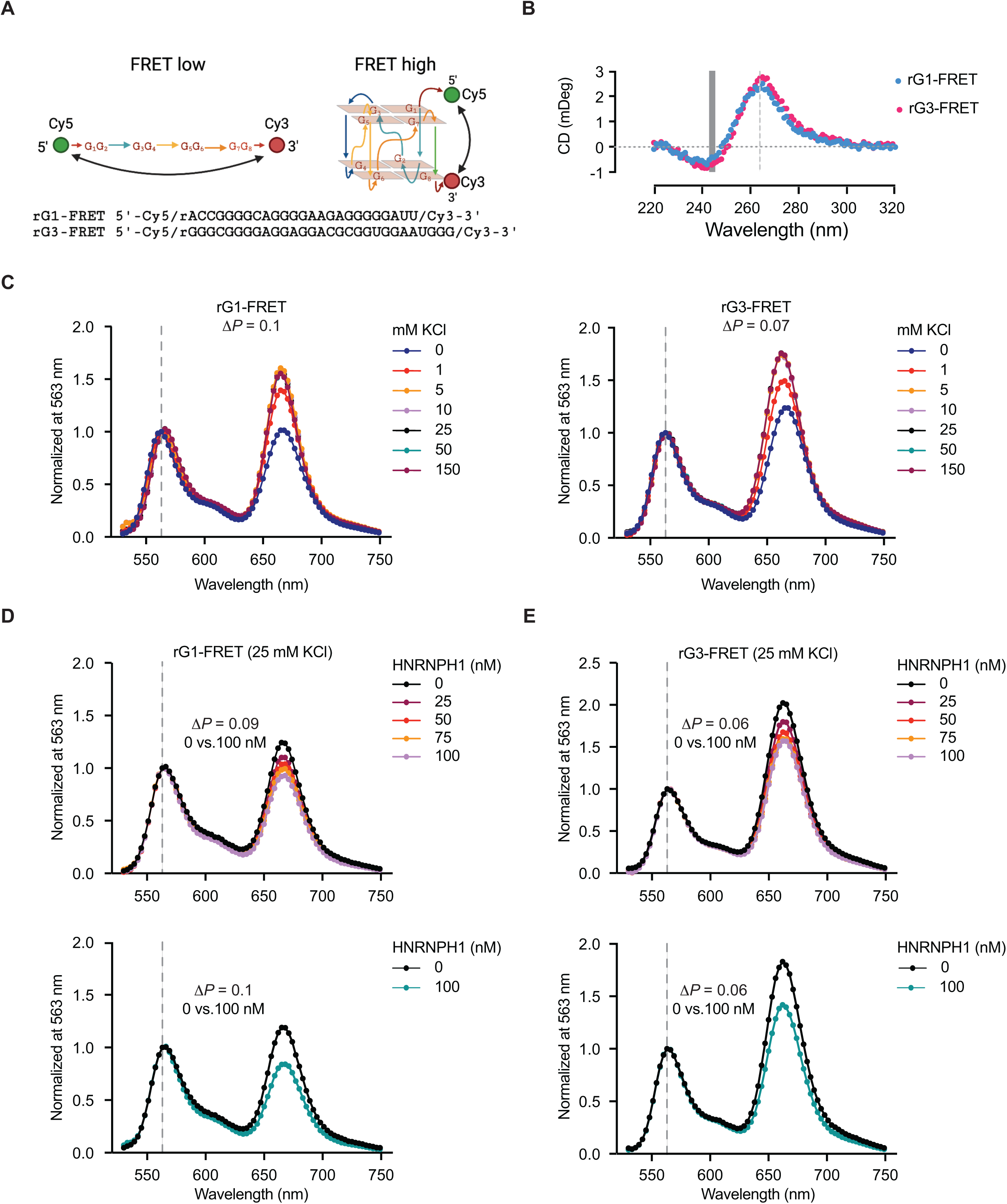
HNRNPH1 destabilizes the G4 structures of the *EWSR1*-exon 8 rG1 and rG3 RNA oligomers. **A.** Schematic of the rG1- and rG3-FRET assays. **B.** CD spectra confirming the folded states rG1- and rG3-FRET RNA oligomers in the presence of 150 nM KCl. **C**. FRET analysis of the rG1 and rG3 RNA oligomers at increasing concentrations of KCl (0 to 150 mM). **D** FRET analysis of the rG1-FRET RNA oligomers at increasing concentrations of HNRNPH1 (0 – 100 nM). The upper and lower panels show results from two independent experiments. **E** FRET analysis of the rG3-FRET RNA oligomers at increasing concentrations of HNRNPH1 (0 – 100 nM). The upper and lower panels show results from two independent experiments. **A** created in BioRender.

To assess the impact of protein on the rG1-FRET and rG3-FRET RNA oligomers at 25 mM KCl, where CD results indicate the rG1 and rG3 oligomers form G4 structures that exhibit thermal melting points of over 50°C, we first determined the effect of the BG4 antibody on the two fluorescent oligomers (**Figure S5B**).Titration of the BG4 antibody yielded minimal changes in the *P* values (rG1-FRET Δ*P*=0.02; rG3-FRET Δ*P*=0.01; 0 versus 100 mM BG4), which, along with the electromobility shift results (**Figure 3A**), shows that BG4 recognizes but does not alter the structure of these RNA oligomers. We next assessed the effect of titrating increased concentrations of HNRNPH1 on the energy transfer of the two FRET oligomers (**Figures 6D** and **E,** and **Table 2**). The addition of 100 nM HNRNPH1 to the rG1-FRET oligomer generated a similar Δ*P* (0.09 or 0.1) as the difference between 0 and 100 mM KCl, consistent with HNRNPH1 destabilizing the folded structure (**Figures 6D**). The addition of HNRNPH1 to the rG3-FRET RNA oligomer also generated a Δ*P* value of 0.06 at 100 nM protein, reflecting a structural transition of a similar magnitude to that seen under different concentrations of KCl (0 and 100 mM; Δ*P*=0.07). These studies suggest that HNRNPH1’s binding of the two-tier G4s formed by rG1 and rG3 can destabilize their structure and induce a non-G4 state.

**Table 2.** FRET efficiency parameters (*P*) for the interaction of HNRNPH1 with *EWSR1*-exon 8 rG1 and rG3 RNA oligomers

| **HNRNPH1 (nM)** | **rG1-FRET** |  |  |  |  | **rG3-FRET** |  |  |  |  |
| --- | --- | --- | --- | --- | --- | --- | --- | --- | --- | --- |
|  | **0\*** | **25** | **50** | **75** | **100** | **0\*** | **25** | **50** | **75** | **100** |
| **Full-length** | 0.45 | 0.49 | 0.51 | 0.52 | 0.54 | 0.37 | 0.39 | 0.41 | 0.42 | 0.43 |
| **qRRM1-2** | 0.45 | 0.47 | 0.50 | 0.58 | 0.62 | 0.37 | 0.39 | 0.40 | 0.41 | 0.42 |
| **qRRM2-3** | 0.46 | 0.50 | 0.50 | 0.50 | 0.51 | 0.35 | 0.37 | 0.37 | 0.37 | 0.38 |

\*Standard error of the mean for RNA oligomers in the absence of protein = 0.02 (rG1 and rG3)

Our cell-based studies indicated that neither HNRNPH2 nor HNRNPF alters the splicing of *EWSR1*-exon 8 even though these proteins are highly homologous (**Figure S6**), and their consensus binding sequences are all G-rich. These non-redundant functions could reflect differences in the recognition and binding of specific sequences or the functional consequences of binding. We thus assessed whether HNRNPH2 or HNRNPF could bind either or both *EWSR1*-exon 8 G-rich sequences using BLI and the FRET assays. Even though the amino sequences of HNRNPH1 and HNRNPH2 differ at just nine residues (five in the qRRM1 domains of each protein, one in the qRRM2 domain, one in the first glycine-rich, and two in the qRRM3 domain), HNRNPH2 showed no binding of either the rG1 or rG3 sequences and no alteration in their respective FRET assays (**Figures S7A** and **B**). Consistent with this observation, BLI analysis of HNRNPH2 indicated minimal binding of rG1 or rG3 (**Figure S7C**). Interestingly, HNRNPF generated a substantial concentration-dependent reduction in the energy transfer from the acceptor to the donor in the case of the rG1-FRET oligomer (Δ*P*=0.25, 0 versus 100 nM protein), which was greater than that observed for HNRNPH1 (**Figure S7D**). In contrast, HNRNPF’s interaction with the rG3-FRET oligomer induced a change in state (Δ*P*= 0.07, 0 versus 100 nM protein), as observed for the interaction of this oligomer with HNRNPH1 or when we compared 0 and 100 nM KCl conditions (**Figure S7E**). This observation is interesting because our cell-based studies have indicated that HNRNPF does not function in the regulation of the exclusion of *EWSR1*-exon 8 and, so we evaluated the hypothesis that the binding affinity and the kinetics of HNRNPF’s interaction with rG1 and rG3 may indicate differences between the interaction of HNRNPH1 and HNRNPF (**Figures S7F**). HNRNPF exhibited sub-nanomolar binding affinities for both rG1 and rG3 (K_D_ = 0.34 and 0.35 nM, respectively), which is stronger than that observed for HNRNPH1’s binding of these sequences (**Figure 4E** and **Table 1**). Analysis of the kinetic parameters suggests that the observed stronger binding is a consequence of a 4-fold slower dissociation for rG1 (HNRNPF, k_d_ = 2.8 E-04 1/s, HNRNPH1 k_d_= 1.1 E-03 1/s) and a 3-fold slower dissociation for rG3 (HNRNPF, k_d_ = 4.4 E-04 1/s, HNRNPH1 k_d_ = 1.5 E-03 1/s). We observed similar binding affinities of HNRNPF to the 7-dG variants at low nanomolar binding affinities (**Figure S7G**) as we observed for HNRNPH1. However, a comparison of the log-log transformation of the k_a_ and k_d_ values obtained from the kinetic fitting of HNRNPF showed it bound faster to the RNA in a G4 state than the 7dG versions, but it exhibited similar dissociation kinetics for RNAs in a G4 or non-G4 state (**Figure S7H**). Collectively, we hypothesize that though HNRNPF can bind both sequences in both states, HNRNPH1’s binding kinetics are critical for generating a functionally relevant interaction.

### The qRRM1-2 domains of HNRNPH1are sufficient to disrupt a G4 structure

The enrichment for differences in the HNRNPH1 and HNRNPH2 qRRM1 and qRRM2 protein domains suggest that these regions are critical determinants of the differences in the BLI and FRET assays. Previous NMR studies showing that the qRRM1 and qRRM2 domains, but not qRRM3, are responsible for recognizing G-rich sequences also indicate the relative importance of these domains in sequence or structure recognition (31, 32). To investigate if the interaction with specific domains of HNRNPH1 contributes to changes in the FRET signals obtained for rG1 and rG3, we used purified proteins corresponding to truncated versions of HNRNPH1, the qRRM1 and qRRM2 domains (qRRM1-2), or the qRRM2, first glycine-rich region and qRRM3 (qRRM2-3) (**Figures 7A**). Using the FRET-RNA oligomers, we observed that the HNRNPH1 qRRM1-2 truncated protein induced the same Δ*P*=0.1 for the rG1-FRET oligomer as the full-length protein, but minimal change in the energy transfer between the donor and acceptor of the rG3-FRET oligomer (**Figures 7B**). We observed no changes in the FRET assays in the presence of the HNRNPH1 qRRM2-3 truncated protein (**Figure 7C**), demonstrating the selectivity and sensitivity of the assay and highlighting the importance of the qRRM1 and qRRM2 domains of the HNRNPF/H proteins described previously (43).

**Figure 7:**
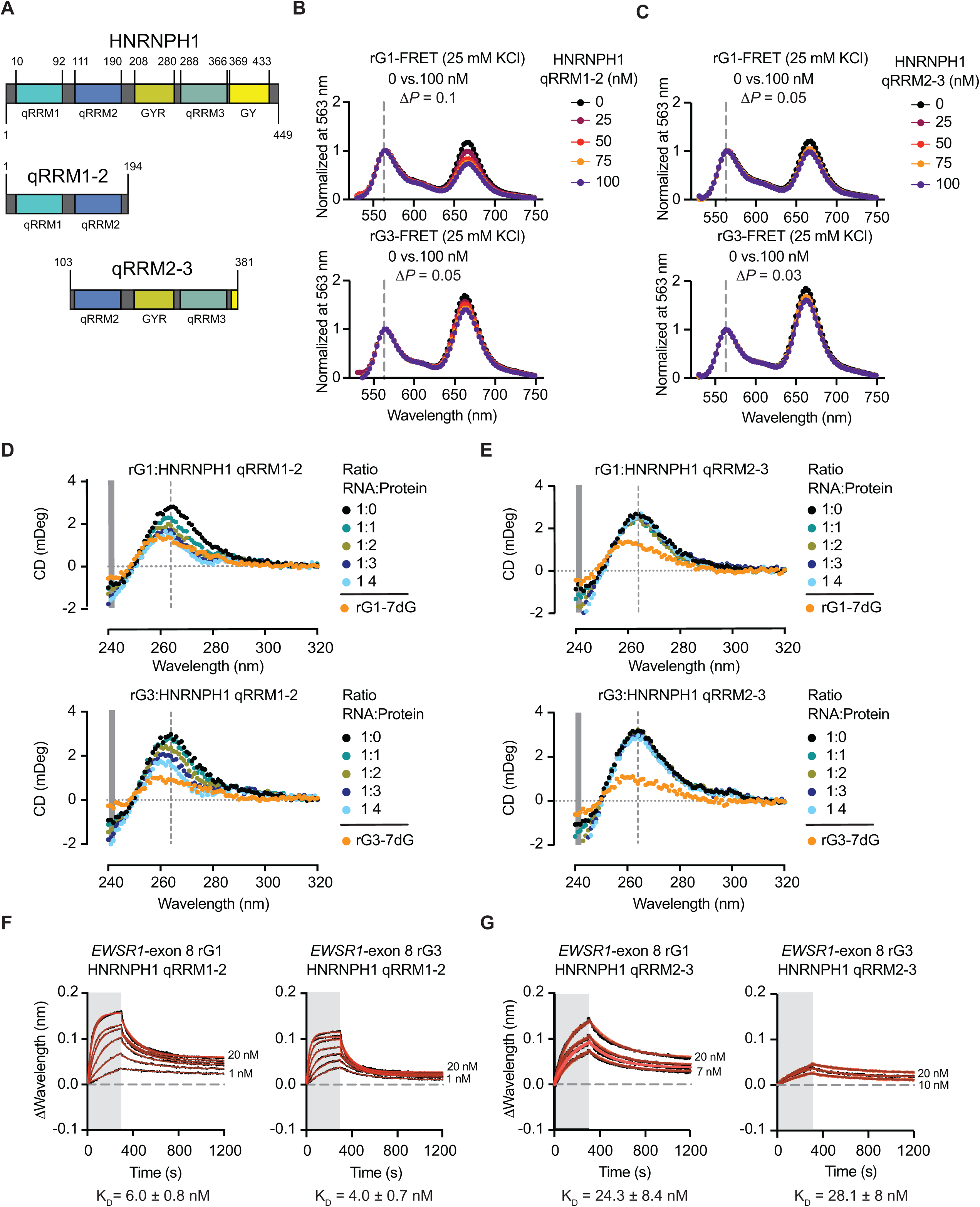
HNRNPH1 domains and RNA structure. **A.** Schematic of full length HNRNPH1 and the qRRM1-2 and qRRM2-3 truncated HNRNPH1 proteins. **B**. FRET analysis of the rG1 and rG3 RNA oligomers at increasing concentrations of truncated HNRNPH1 proteins, qRRM1-2, (0 – 100 nM). **C.** FRET analysis of the rG1 and rG3 RNA oligomers at increasing concentrations of truncated HNRNPH1 protein, qRRM2-3 (0 – 100 nM). **D.** CD spectra for the indicated RNA oligomers in the presence of increasing concentrations (0 – 6 µM) the qRRM1-2 truncated HNRNPH1 protein. **E.** CD spectra for the indicated RNA oligomers in the presence of increasing concentrations (0 – 6 µM) the qRRM2-3 truncated HNRNPH1 protein. **F.** Sensorgrams for the indicated RNA oligomers and the qRRM1-2 truncated HNRNPH1 protein. **G.** Sensorgrams for the indicated RNA oligomers and the qRRM2-3 truncated HNRNPH1 protein.

The truncated HNRNPH1 domains are less subject to aggregation at higher concentrations (µM) than the full-length proteins, allowing us to next employ these in CD assays as an orthogonal method to assess the effect of HNRNPH1’s interaction on the structure of a folded RNA. First, we confirmed that each protein generated an expected spectrum – no negative peak at 240 nm and no positive peak at 264 nm (**Figure S8A**), suggesting that we could use changes in the CD spectra at the latter wavelength to determine if protein binding shifts the CD typical of rG4s to that resembling a different structure. The addition of the truncated HNRNPH1 qRRM1-2 generated a substantial reduction in the 264 nm ellipticity to similar CD spectra of rG1-7dG and rG3-7dG at 4:1 protein to RNA ratio (**Figure 7D**). In contrast, we observed little to no change in the RNA CD spectra in the presence of the qRRM2-3 truncated protein (**Figure 7E**). These results suggest that the HNRNPH1 qRRM1-2 domains are critical for interacting with and altering the G4 state of the rG1 and rG3 RNA oligomers. To confirm that the HNRNPH1 qRRM1-2 and qRRM2-3 domains exhibit different affinities for the rG1 and rG3 RNA oligomers, we performed BLI analysis using each of these truncated proteins (**Figure 7F**and **G**). The qRRM1-2 protein exhibited low nM binding affinities for rG1 (6 nM) and rG3 (4 nM), consistent with these domains contributing a substantial proportion of HNRNPH1’s affinity for these RNA oligomers, particularly as the qRRM2-3 proteins exhibited a five-fold weaker binding of these oligomers. The kinetic binding parameters indicated different association rates for the interactions between the qRRM1-2 and the qRRM2-3 domains, though comparable dissociation rates (**Figure S8B**). Together, the FRET and CD results suggest that the qRRM1-2 domains are crucial for recognizing and generating the optimal protein-RNA complexes involving one or more G-rich sequences that contribute to an HNRNPH1-dependent splicing event.

## DISCUSSION

The detection of many thousands of G-rich sequences within the human transcriptome that, under physiological conditions, can form G-quadruplexes has led to an extensive discussion of whether the function of some RNA-binding proteins is to counter the formation of these complex structures transcriptome-wide, or whether at least some of these rG4-RNA-binding protein interactions are dynamic and regulate gene expression (14,22,69). Studies of the HNRNPH/F protein family have indicated that their interaction with G-rich sequences can alter RNA folding, and this may have transcriptome-wide effects, though these proteins also regulate specific RNA processing events. However, limitations of some of these studies include the extrapolation of findings generated using one HNRNPH/F protein to all members of this protein family and the use of artificial RNA G-rich sequences to define structure-function relationships. Here, we have used a model system based on a disease-relevant HNRNPH1-dependent splicing event to enhance our understanding of its function as a splicing factor and to probe the biophysical properties of its interaction with disease-relevant G-rich sequences. Our results show that HNRNPH1’s binding of a two-tier quartet G-quadruplex destabilizes this secondary structure in a non-catalytic fashion to generate an RNA with properties consistent with that of a non-G4 state. Moreover, the kinetics of HNRNPH1’s binding favors the accumulation of RNA in a non-G4 state.

Understanding the biological functions of the HNRNPH/F proteins is critical, given the importance of these proteins in regulating RNA metabolism. Here, we demonstrated that study of the HNRNPH1-dependent processing of *EWSR1*-exon 8 has disease relevance and warrants further study by determining that a substantial proportion of Ewing sarcomas diagnosed each year will depend on the efficient exclusion of *EWSR1*-exon 8 to express the fusion oncoprotein that initiates and promotes tumorigenesis. We speculate that the exclusion of *EWSR1*-exon 8 in the context of the *EWS-FLI1* transcript occurs as an adaption of an existing HNRNPH1-mediated splicing event, and our long-read sequencing also highlighted ways in which HNRNPH1 function may regulate the expression of *EWSR1*. The *EWSR1* gene expresses many protein-coding and non-coding transcripts. Based on data reported for multiple tissue types (GTEx) of the protein-coding *EWSR1* transcripts, *EWSR1-207* (ENST00000406548) is the predominantly expressed variant (median TPM 41.8; range 12.5 – 94.4), while the expression of *EWSR1-206* (ENST00000397938) is lower (median TPM 1.015; range 1.015 – 9.59). Our long-read sequencing data combined with coding-potential analysis helped us identify transcripts that align with *EWSR1-206* and *EWSR1-207* as expressed at similar levels in TC-32 cells. This same combination of analyses showed a loss of *EWSR1-207* transcripts and dominance of *EWSR1-206* in the absence of HNRNPH1. One explanation for this observation may be the predicted strengths of the 3’ splice site for *EWSR1*-intron 8 used to generate the two forms of *EWSR1*-Exon 9 (38 versus 35 nucleotides) that distinguish *EWSR1-206 and EWSR1-207.* The strength of the *EWSR1-206* 3’ splice site is estimated at 7.90, whereas the *EWSR1-207* 3’ splice site has a strength of 5.26. These differences suggest that HNRNPH1 may facilitate the use of the weaker 3’ splice site, a hypothesis we will explore in future studies. Critically, our observation that the depletion of HNRNPH1 results in the continued expression of an *EWSR1* protein-coding transcript. This observation suggests that it should be feasible to block HNRNPH1’s interaction with a specific *cis*-acting sequence that regulates EWS-FLI1 expression while maintaining EWSR1 protein expression. Interestingly, though our minigene constructs proved valuable in establishing the selectivity of HNRNPH1 function in mediating *EWSR1-*exon 8 exclusion, neither fully recapitulate the results of our long-read sequencing, which showed complete inclusion of *EWSR1*-exon 8 in the context of *EWSR1* transcripts, and almost complete exclusion in the context of the fusion transcript. This observation suggests that other *cis*-acting regulatory elements outside of those incorporated into the two minigenes contribute to *EWSR1*-exon 8 inclusion and exclusion, and future studies will aim to identify these sequences and whether HNRNPH1 or another RBP bind them.

The modified minigenes allowed us also to assess the effect of disrupting the longest G-tracts present in the 3’ end of *EWSR1*-exon 8. Disruption of the three G-tracts within the 106 – 124 nt region of *EWSR1*-exon 8 inclusion indicates that this sequence can function as a *cis*-regulatory element defining the splicing of *EWSR1*-exon 8. Furthermore, our biophysical studies show that HNRNPH1 can bind this sequence in a non-G4 and G4 state and that if in a G4 state, HNRNPH1 binding could destabilize this secondary structure. Collectively, these results indicate that targeting this 106 – 124 nt *cis*-regulatory element through, for example, the use of antisense oligonucleotides warrants study as a means of disrupting EWS-FLI1 expression. However, the complexity of the G-rich region closer to the 3’ end of *EWSR1*-exon 8 (150 – 181 nts) will require further study to define the minimal *cis*-regulatory element present in this region. Particularly intriguing is our finding that three G-A substitutions towards the 3’ end of the exon (nts 166, 174, and 181) resulted in *EWSR1-*exon 8 exclusion, but the same nucleotide changes in combination with others within the upstream region resulted in exon retention. We evaluated the effect of adding further G-As because this eliminated the localized formation of any quadruplex structure. However, determining the specific contribution of the eight G-tracts within this region to the splicing of *EWSR1*-exon 8 will require additional study. Our biophysical studies focused on the rG3 RNA oligomer because of its contrasting properties with those of the rG1 RNA oligomer. Future studies should also examine whether slight variations in the sequence, such as examining the rG2 sequence in more detail or RNA oligomers encompassing the entire 150 – 181 nt region alter these properties.

G-rich RNAs can only form parallel G-quadruplex structures, and though most studies of RNA G4s have focused on folds that stack three G-tetrads, there is evidence that RNA can form non-canonical G4s that include stacks of two G-tetrads and G4s with different loop lengths. We previously reported enrichment for a GG-N_7_-GG-N_7_-GG-N_7_-GG-N_7_ motif at exonic sites reported as bound by HNRNPH1 binding sites (23). This sequence motif is compatible with forming a two-layer G-quartet structure that a study by Zhang and Balasubramanian (2012) predicted would have thermodynamic properties more compatible with an active regulatory function than canonical three-layer G4s (46). While both the CD spectra of oligomers corresponding to the rG1 and rG3 *EWSR1*-exon 8 sequences demonstrate that these oligomers form parallel G4s and modeling predicts these sequences form two-tier G quartets, we noted several differences in the thermodynamic characteristics of the two RNA oligomers, related to their sequence composition and length (25 versus 29 nts). For example, we observed that rG3 exhibited a CD spectrum consistent with the formation of a G4 structure in the presence of as little as 1 mM KCl and had a higher T_m_ than rG1 (68°C versus 50°C). Multiple factors can contribute to these differences, including which guanine residues form the planar tetrad and thus the loop composition and length of each oligomer. For example, previous studies have shown that in the case of an artificial sequence that formed a two-tier G4 structure, the longer the loop length, the lower the T_m_ (21). Here, modeling the endogenous human sequences that correspond to rG1 and rG3, we observe that the rG1 sequence will form a two-tier G4 with an internal G-tetrad that contains uneven loops of 1 to 5 nts, while the rG3 could form various quadruplexes with loop lengths and conformations. Considering previous findings correlating loop length and thermostability, we predict that rG3 will form G4 structures that contain loops of 1 to 3 nts. We speculate that these different conformations – a two-tier G4 with an internal G-tetrad that contains uneven loops of 1 to 5 nts versus G4 structures that contain loops of 1 to 3 nts may contribute to the lower thermal melting point we observed for rG1 versus rG3. Both rG1 and rG3 exhibit strong dependence on changes in bulk salt concentration, indicating the G4 formation process is cooperative. Regardless, we observed higher thermal melting values for these G-tracts compared to synthetic two-tier G4 reported (20, 21). Defining the exact lengths of the loops with the G4 structures that endogenous G-rich sequences such as rG1 and rG3 could form and the effect of the nucleotide composition of any loop will require further study. Critically, such differences in the thermostability of G-rich RNA regions could be of biological relevance, as the capacity of an RNA to fold into G4 structures at different rates, as well as the effect of RNA binding proteins on the folding, destabilization, or resolution of such structures has the potential to alter multiple aspects of RNA metabolism, including alternative splicing.

The homology of the HNRNPH/F protein family suggests their functional differences depend on the amino acid compositions of their qRRM domains and variations in the second glycine-rich region. Consistent with our previous work and the results of our cell-based minigene studies that showed silencing of *HNRNPH2* has no effect on the relative inclusion or exclusion of *EWSR1*-exon 8, we observed that HNRNPH2 did not alter either the rG1 or rG3-FRET signals or bind the *EWSR1*-exon 8 G-rich sequences. Interestingly, the results for HNRNPF generated contrasting data. Based on our previous results, as expected, we observed that depletion of HNRNPF did not affect the splicing of *EWSR1*-exon 8; however, FRET and BLI studies showed that the HNRNPF can bind both rG1 and rG3 sequences with low nM affinities and can alter their structures. Previous studies showed that HNRNPH/F have differential qRRM inter-dynamics resulting in HNRNPH1-RNA complexes adopting a compact conformation and HNRNPF-RNA complexes forming a more extended conformation (44). Future studies will probe whether these differences in conformation are critical to the destabilization of G4 RNA structures observed in this study and their functions in RNA processing. Our results also suggest that while both HNRNPH1 and HNRNPF can alter the structure of an RNA in a cell-free system, our cell-based experiments indicate that this is insufficient to mediate the exclusion of EWSR1 exon 8. Determining whether this reflects differences in HNRNPH1’s interaction with additional cis-regulatory sequences or other splicing factors from that of HNRNPF could prove critical for understanding the respective functions of these two closely related RNA binding proteins and warrants additional study.

Using BLI analysis, we demonstrate distinct kinetic differences for the binding of HNRNPH1 with rG1 and rG3 oligomer depending on whether the RNA is in a non-G4 or G4 state. Specifically, HNRNPH1’s association rate kinetic is three times faster for rG1 than rG1-7dG and ten times faster for rG3 than rG3-7dG. Assuming that during synthesis of the nascent RNA containing these sequences, there is an equilibrium between the rate at which these G-rich regions remain in unfolded structure or fold into G4s, these kinetic parameters suggest that HNRNPH1’s dominant interaction will be with RNA in a G4 state. However, once bound, our FRET studies indicate that HNRNPH1 can destabilize the two-tier G4s formed by rG1 and rG3 in that we observed a change in the FRET signals at 0 versus 100 nM HNRNPH1 resembling that observed when we compared the signals in the presence and absence of KCl. Supporting these results, we observe that a truncated HNRNPH1 qRRM1-2 protein generates a change in the CD spectra in both oligomers that an RNA: Protein ratio of 1:4 resembles the spectra of the respective 7dG variants. Critically, the addition of protein alone does not exert this effect as a truncated HNRNPH1 qRRM2-3 protein-induced no change in the CD spectra of either RNA oligomer. Of note, both the FRET and CD studies suggest that HNRNPH1 has a more significant destabilizing effect on rG1 than rG3, which potentially relates to the differences in the thermal stability of the two RNAs discussed above.

In conclusion, this study shows that HNRNPH1 can induce a non-catalytic destabilization of the secondary G4 structures formed by G-rich sequences. Significantly, these sequences contribute to an exon exclusion event necessary for expressing the fusion oncoprotein in a subset of Ewing sarcoma. Further study of HNRNPH1’s regulation of this exon exclusion event will offer opportunities to inhibit it selectively and enhance our understanding of how HNRNPH1’s interaction with G-rich sequences alters the secondary structure of RNA to regulate RNA processing.

## DATA AVAILABLITY

Full size images related to **Figure 4C** are included as a supplementary data file. The minigene plasmids are available upon reasonable request. RNA sequencing datasets have been deposited in the SRA repository (project ID: PRJNA779048).

## SUPPLEMENTARY INFORMATION

Supplementary data available.

## ACKNOWLEDGEMENT

We thank Grzegorz Piszczek and Di Wu, Biophysics Core Facility, National Heart, Lung, and Blood Institute, NIH for their assistance with this project. We thank Caroline Fromont and other members if the CCR Sequencing facility, as well as Yongmei Zhao, Xiongfeng Chen, and Xinyu Chen (CCR-SF Bioinformatics Group (CCR-SF IFX), Biomedical Informatics and Data Science (BIDS), Frederick National Laboratory for Cancer Research (FNLCR) for the performance of the long-read sequencing and their assistance with the analysis of these data. We also thank Alexi Lobanov and Mayank Tandon (CCR Collaborative Bioinformatics Resource) and Hsien-Chao Chou (Oncogenomics Section, Genetics Branch, CCR, NCI) for useful discussion of the *EWSR1* and *EWS-FLI1* long-read sequencing analysis. We also thank members of the Functional Genetics Section, Genetics Branch for their constructive discussion of this project and careful reading of the manuscript. The CCR Genomics Core, CCR, NCI performed the Sanger sequencing.

