## Supplementary Information for "HNRNPH1 destabilizes the G-quadruplex structures formed by G-rich RNA sequences that regulate the alternative splicing of an oncogenic fusion transcript"

**Supplementary Tables**

**Table S1:** Long-read sequencing of *EWS-FLI1* (TC-32 cells transfected with the indicated siRNAs)

**Table S2:** Long-read sequencing of *EWSR1* transcripts (TC-32 cells transfected with the indicated siRNAs)

**Supplementary Figures**

**Figures S1- S8**

**Supplementary data:** Full size images related to **Figure 4C**

**Table S1: Long-read sequencing of *EWS-FLI1* (TC-32 cells transfected with the indicated siRNAs)**

| siNEG - *EWS-FLI1* | | | | | | | | | | |
|---|---|---|---|---|---|---|---|---|---|---|
| **IsoSeq ID** | **Chromosome 11 coordinates** | **Chromosome 22 coordinates** | **Transcript ID** | **Transcript length** | **Peptide length** | **Fickett score** | **isoelectric point (pI)** | **ORF integrity** | **Coding probability** | **Comments** |
| PBfusion.458 | chr11:128805365-128812415 (+) | chr22:29267961-29287134 (+) | transcript/119484 | 3303 | 499 | 0.35 | 8.46 | 1 | 1.0 | Full length 7/6 *EWS-FLI1* |
| PBfusion.463 | chr11:128805365-128813265 (+) | chr22:29268292-29287134 (+) | transcript/92861 | 3818 | 499 | 0.34 | 8.46 | 1 | 1.0 | Full length 7/6 *EWS-FLI1* |
| PBfusion.459 | chr11:128805364-128812415 (+) | chr22:29268281-29288786 (+) | transcript/124789 | 3160 | 329 | 0.36 | 6.54 | 1 | 1.0 | *EWSR1* Exon 8 inclusion |
| PBfusion.461 | chr11:128803887-128812415 (+) | chr22:29268292-29287134 (+) | transcript/130691 | 3037 | 522 | 0.36 | 9.01 | 1 | 1.0 | Additional *FLI1* sequences |
| PBfusion.460 | chr11:128805365-128812415 (+) | chr22:29268267-29287134 (+) | transcript/141333 | 2826 | 443 | 0.35 | 8.50 | 1 | 1.0 | *EWSR1* Exon 6 exclusion |
| PBfusion.455 | chr11:128782310-128786066 (+) | chr22:29268292-29355244 (+) | transcript/7649 | 6816 | 274 | 0.32 | 4.46 | 1 | 1.0 | Truncated; retained intron |
| PBfusion.456 | chr11:128782310-128783884 (+) | chr22:29268269-29355244 (+) | transcript/53850 | 4681 | 274 | 0.35 | 4.46 | 1 | 1.0 | Truncated; retained intron |
| PBfusion.457 | chr11:128782310-128783412 (+) | chr22:29268292-29355244 (+) | transcript/76113 | 4186 | 274 | 0.36 | 4.46 | 1 | 1.0 | Truncated; retained intron |
| PBfusion.462 | chr11:128805365-128811923 (+) | chr22:29268282-29287134 (+) | transcript/72376 | 4289 | 207 | 0.28 | 7.79 | 1 | 0.9 | Truncated; retained intron |

| siHNRNPH1 - *EWS-FLI1* | | | | | | | | | | |
|---|---|---|---|---|---|---|---|---|---|---|
| **IsoSeq ID** | **Chromosome 11 coordinates** | **Chromosome 22 coordinates** | **Transcript ID** | **Transcript length** | **Peptide_length** | **Fickett_score** | **isoelectric point (pI)** | **ORF_integrity** | **Coding probability** | **Comments** |
| PBfusion.380 | chr11:128805365-128812415 (+) | chr22:29268281-29287134 (+) | transcript/102452 | 2979 | 499 | 0.36 | 8.46 | 1 | 1.0 | Full length 7/6 *EWS-FLI1* |
| PBfusion.379 | chr11:128805364-128812415 (+) | chr22:29268295-29288786 (+) | transcript/97074 | 3146 | 329 | 0.36 | 6.54 | 1 | 1.0 | *EWSR1*-Exon 8 inclusion |
| PBfusion.378 | chr11:128805364-128812415 (+) | chr22:29268310-29288786 (+) | transcript/37400 | 5048 | 329 | 0.32 | 6.54 | 1 | 1.0 | Truncated; retained intron |
| PBfusion.377 | chr11:128782310-128786029 (+) | chr22:29268298-29355244 (+) | transcript/9075 | 6771 | 274 | 0.32 | 4.46 | 1 | 1.0 | Truncated; retained intron |

| siFLI1 - *EWS-FLI1* | | | | | | | | | | |
|---|---|---|---|---|---|---|---|---|---|---|
| **IsoSeq ID** | **Chromosome 11 coordinates** | **Chromosome 22 coordinates** | **Transcript ID** | **Transcript length** | **Peptide_length** | **Fickett_score** | **isoelectric point (pI)** | **ORF_integrity** | **Coding probability** | **Comments** |
| PBfusion.353 | chr11:128805365-128812415 (+) | chr22:29268293-29287134 (+) | transcript/123171 | 2967 | 499 | 0.36 | 8.46 | 1 | 1.0 | Full length 7/6 *EWS-FLI1* |
| PBfusion.351 | chr11:128782310-128783892 (+) | chr22:29268292-29355244 (+) | transcript/52565 | 4666 | 274 | 0.35 | 4.46 | 1 | 1.0 | Truncated; retained intron |
| PBfusion.352 | chr11:128782480-128783892 (+) | chr22:29268281-29287419 (+) | transcript/141279 | 2537 | 274 | 0.39 | 4.46 | 1 | 1.0 | Truncated; retained intron |
| PBfusion.350 | chr11:128782310-128786029 (+) | chr22:29268015-29355244 (+) | transcript/6760 | 7059 | 246 | 0.29 | 9.49 | 1 | 0.9 | Truncated; retained intron |

### Table S2: Long-read sequencing of *EWSR1* transcripts (TC-32 cells transfected with the indicated siRNAs)

**siNeg - *EWSR1***

| IsoSeq ID | chromosome coordinates | Transcript ID | Comments | Transcript length | 5'UTR and exon 1 (nts) | Exon 2 (nts) | Exon 3 (nts) | Exon 4 (nts) | Exon 5 (nts) | Exon 6 (nts) | Exon 7 (nts) | Exon 8 (nts) | Exon 9 (nts) | Exon 10 (nts) | Exon 11 (nts) | Exon 12 (nts) | Exon 13 (nts) | Exon 14 (nts) | Exon 15 (nts) | Exon 16 (nts) | Exon 17 and 3'UTR (nts) | peptide length | Fickett score | isoelectric point (pI) | ORF integrity | Coding probability |
|---|---|---|---|---|---|---|---|---|---|---|---|---|---|---|---|---|---|---|---|---|---|---|---|---|---|---|
| **PB.15543.1** | **chr22:29268037-29300516 (+)** | **transcript/149741** | ***closest match: EWSR1-207 ENST00000406548*** | **2625** | **313** | **37** | **52** | **124** | **187** | **168** | **212** | **181** | **35** | **33** | **119** | **130** | **123** | **163** | **98** | **253** | **395** | **656** | **0.42** | **9.37** | **1** | **1.0** |
| **PB.15543.44** | **chr22:29268299-29300334 (+)** | **transcript/163308** | ***closest match: EWSR1-206 ENST00000397938*** | **2182** | **51** | **37** | **52** | **124** | **187** | **168** | **212** | **181** | **38** | **33** | **119** | **130** | **123** | **163** | **98** | **253** | **214** | **657** | **0.43** | **9.37** | **1** | **1.0** |
| PB.15543.8 | chr22:29268268-29300512 (+) | transcript/161890 | *closest match: EWSR1-203 ENST00000332050* | 2285 | 82 | 37 | 52 | 124 | 187 | 168 | 212 | Alt Exon: 76nt; 200nt ds | 35 | 33 | 119 | 130 | 123 | 163 | 98 | 253 | 391 | 621 | 0.42 | 9.28 | 1 | 1.0 |
| PB.15543.30 | chr22:29268290-29300516 (+) | transcript/163868 | closest match: *EWSR1-202* ENST00000332035 | 2208 | 60 | 37 | 52 | 124 | 187 | Excluded | 212 | 181 | 38 | 33 | 119 | 130 | 123 | 163 | 98 | 253 | 395 | 573 | 0.45 | 9.78 | 1 | 1.0 |
| PB.15543.9 | chr22:29268268-29300334 (+) | transcript/162761 | | 2254 | 82 | 37 | 52 | 124 | 187 | 168 | 212 | 181 | 38 | 33 | 119 | 130 | 123 | 163 | Alt Exon: 137 nt; alt 5' splice | 253 | 214 | 670 | 0.43 | 9.24 | 1 | 1.0 |
| PB.15543.15 | chr22:29268270-29300334 (+) | transcript/165040 | | 2175 | 80 | 37 | 52 | 124 | 187 | 168 | 212 | 181 | Excluded | 33 | 119 | 130 | 123 | 163 | 98 | 253 | 214 | 329 | 0.42 | 6.70 | 1 | 1.0 |
| PB.15543.21 | chr22:29268282-29300516 (+) | transcript/157384 | | 2423 | 68 | 37 | 52 | 124 | 187 | 168 | 212 | 181 | 38 | 33 | 119 | 130 | 123 | 163 | Alt Exon: 137 nt; alt 5' splice | 253 | 398; TT insertion | 670 | 0.43 | 9.24 | 1 | 1.0 |
| PB.15543.22 | chr22:29268282-29300512 (+) | transcript/162190 | | 2252 | 68 | 37 | 52 | 124 | 187 | 168 | 212 | 181 | 35 | 33 | 119 | 130 | Excluded | 163 | 98 | 253 | 391 | 615 | 0.42 | 8.96 | 1 | 1.0 |
| PB.15543.26 | chr22:29268288-29300335 (+) | transcript/163981 | | 2182 | 62 | 37 | 52 | 124 | 187 | 168 | 212 | 181 | 38 | 33 | 119 | 130 | 123 | 163 | Alt Exon: 86 nt; alt 5' splice and 3' splice | 253 | 215 | 653 | 0.43 | 9.20 | 1 | 1.0 |
| PB.15543.27 | chr22:29268289-29300515 (+) | transcript/160103 | | 2338 | 61 | 37 | 52 | 124 | 187 | 168 | 212 | 181 | Excluded | 33 | 119 | 130 | 123 | 163 | 98 | 253 | 397; TT insertion | 329 | 0.42 | 6.70 | 1 | 1.0 |
| PB.15543.36 | chr22:29268293-29300524 (+) | transcript/160580 | | 2332 | 57 | 37 | 52 | 124 | 187 | 168 | 212 | 181 | 38 | 33 | 119 | 130 | 123 | 163 | Alt Exon: 47 nt; alt 3' splice | 253 | 406: TTT insertion | 640 | 0.43 | 9.34 | 1 | 1.0 |

**siHNRNPH1 - *EWSR1***

| IsoSeq ID | chromosome coordinates | Transcript ID | Comments | Transcript length | 5'UTR and exon 1 (nts) | Exon 2 (nts) | Exon 3 (nts) | Exon 4 (nts) | Exon 5 (nts) | Exon 6 (nts) | Exon 7 (nts) | Exon 8 (nts) | Exon 9 (nts) | Exon 10 (nts) | Exon 11 (nts) | Exon 12 (nts) | Exon 13 (nts) | Exon 14 (nts) | Exon 15 (nts) | Exon 16 (nts) | Exon 17 and 3'UTR (nts) | peptide length | Fickett score | isoelectric point (pI) | ORF integrity | Coding probability |
|---|---|---|---|---|---|---|---|---|---|---|---|---|---|---|---|---|---|---|---|---|---|---|---|---|---|---|
| **PB.13348.21** | **chr22:29268293-29300515 (+)** | **transcript/116945** | **closest match: *EWSR1-206 ENST00000397938*** | **2369** | **57** | **37** | **52** | **124** | **187** | **168** | **212** | **181** | **38** | **33** | **119** | **130** | **123** | **163** | **98** | **253** | **395** | **657** | **0.41966** | **9.37005615** | **1** | **1.0** |
| **PB.13348.23** | **chr22:29268293-29300334 (+)** | **transcript/120339** | **closest match: *EWSR1-206* ENST00000397938** | **2188** | **57** | **37** | **52** | **124** | **187** | **168** | **212** | **181** | **38** | **33** | **119** | **130** | **123** | **163** | **98** | **253** | **214** | **657** | **0.42784** | **9.37005615** | **1** | **1.0** |
| PB.13348.8 | chr22:29268270-29300515 (+) | transcript/120388 | *closest match: EWSR1-202 ENST00000332035* | 2227 | 80 | 37 | 52 | 124 | 187 | Excluded | 212 | 181 | 38 | 33 | 119 | 130 | 123 | 163 | 98 | 253 | 397; TT insetion | 601 | 0.42786 | 9.43011475 | 1 | 1.0 |
| PB.13348.5 | chr22:29268268-29300515 (+) | transcript/118572 | | 2331 | 82 | 37 | 52 | Alt exon:127 nt; alt 5' splice | 187 | 168 | 212 | Alt exon: 150 nt; alt 3' splice | Excluded | 33 | 119 | 130 | 123 | 163 | 98 | 253 | 395 | 635 | 0.42042 | 9.24139404 | 1 | 1.0 |
| PB.13348.4 | chr22:29268268-29300515 (+) | transcript/116632 | | 2433 | 82 | 37 | 52 | 124 | 187 | 168 | 212 | 181 | 35 | 33 | 119 | 130 | 123 | 163 | Alt Exon: 137 nt; alt 5' splice | 253 | 395 | 669 | 0.41966 | 9.23846436 | 1 | 1.0 |
| PB.13348.6 | chr22:29268268-29300334 (+) | transcript/119970 | | 2250 | 82 | 37 | 52 | 124 | 187 | 168 | 212 | 181 | 35 | 33 | 119 | 130 | 123 | 163 | Alt Exon: 137 nt; alt 5' splice | 253 | 214 | 669 | 0.42619 | 9.23846436 | 1 | 1.0 |
| PB.13348.9 | chr22:29268270-29300334 (+) | transcript/121271 | | 2174 | 80 | 37 | 52 | 124 | 187 | 168 | 212 | 181 | Excluded | 33 | 119 | 130 | 123 | 163 | 98 | 253 | 214 | 329 | 0.42036 | 6.70159912 | 1 | 1.0 |
| PB.13348.14 | chr22:29268288-29300516 (+) | transcript/118222 | | 2337 | 62 | 37 | 52 | 124 | 187 | 168 | 212 | 181 | Excluded | 33 | 119 | 130 | 123 | 163 | 98 | 253 | 396 | 329 | 0.42716 | 6.70159912 | 1 | 1.0 |
| PB.13348.22 | chr22:29268293-29300515 (+) | transcript/118875 | | 2321 | 57 | 37 | 52 | 124 | 187 | 168 | 212 | 181 | 38 | 33 | 119 | 130 | 123 | 163 | Alt Exon: 47 nt; alt 3' splice | 253 | 395 | 640 | 0.41966 | 9.34295654 | 1 | 1.0 |

**siFLI1 - *EWSR1***

| IsoSeq ID | chromosome coordinates | Transcript ID | Comments | Transcript length | 5'UTR and exon 1 (nts) | Exon 2 (nts) | Exon 3 (nts) | Exon 4 (nts) | Exon 5 (nts) | Exon 6 (nts) | Exon 7 (nts) | Exon 8 (nts) | Exon 9 (nts) | Exon 10 (nts) | Exon 11 (nts) | Exon 12 (nts) | Exon 13 (nts) | Exon 14 (nts) | Exon 15 (nts) | Exon 16 (nts) | Exon 17 and 3'UTR (nts) | peptide length | Fickett score | isoelectric point (pI) | ORF integrity | Coding probability |
|---|---|---|---|---|---|---|---|---|---|---|---|---|---|---|---|---|---|---|---|---|---|---|---|---|---|---|
| **PB.14774.1** | **chr22:29267996-29300335 (+)** | **transcript/142871** | **closest match: *EWSR1-206 ENST00000397938*** | **2487** | **340** | **37** | **52** | **124** | **187** | **168** | **212** | **181** | **38** | **33** | **119** | **130** | **123** | **163** | **98** | **253** | **214** | **657** | **0.41964** | **9.37005615** | **1** | **1.0** |
| **PB.14774.2** | **chr22:29268033-29300516 (+)** | **transcript/137668** | **closest match: *EWSR1-206* ENST00000397938** | **2632** | **317** | **37** | **52** | **124** | **187** | **168** | **212** | **181** | **38** | **33** | **119** | **130** | **123** | **163** | **98** | **253** | **395** | **644** | **0.38912** | **9.50640869** | **1** | **1.0** |
| PB.14774.5 | chr22:29268268-29300516 (+) | transcript/150595 | | 2226 | 82 | 37 | 52 | 124 | 187 | Excluded | 212 | 181 | 35 | 33 | 119 | 130 | 123 | 163 | 98 | 253 | 397; TT insertion | 600 | 0.42786 | 9.43011475 | 1 | 1.0 |
| PB.14774.4 | chr22:29268268-29300516 (+) | transcript/147095 | | 2358 | 82 | 37 | 52 | 124 | 187 | 168 | 212 | 181 | Excluded | 33 | 119 | 130 | 123 | 163 | 98 | 253 | 395 | 329 | 0.41847 | 6.70159912 | 1 | 1.0 |
| PB.14774.7 | chr22:29268280-29300334 (+) | transcript/149701 | | 2241 | 70 | 37 | 52 | 124 | 187 | 168 | 212 | 181 | 38 | 33 | 119 | 130 | 123 | 163 | Alt Exon: 137 nt; alt 5' splice | 253 | 213 | 670 | 0.42619 | 9.23846436 | 1 | 1.0 |
| PB.14774.11 | chr22:29268282-29300334 (+) | transcript/152802 | | 2150 | 68 | 37 | 52 | 124 | 187 | 168 | 212 | 181 | 38 | 33 | 119 | 130 | 123 | 163 | Alt Exon: 47 nt; alt 3' splice | 253 | 213 | 640 | 0.42784 | 9.34295654 | 1 | 1.0 |
| PB.14774.20 | chr22:29268293-29300334 (+) | transcript/155845 | | 2018 | 57 | 37 | 52 | 124 | 187 | Excluded | 212 | 181 | Alt exon: 36 nt; 2 nt deletion | 33 | 119 | 130 | 123 | 163 | 98 | 253 | 213 | 318 | 0.44629 | 9.93341064 | 1 | 1.0 |
| PB.14774.27 | chr22:29268311-29300512 (+) | transcript/144741 | | 2388 | 39 | 37 | 52 | 124 | 187 | 168 | 212 | 181 | 38 | 33 | 119 | 130 | 123 | 163 | Alt Exon: 137 nt; alt 5' splice | 253 | 391 | 670 | 0.41966 | 9.23846436 | 1 | 1.0 |

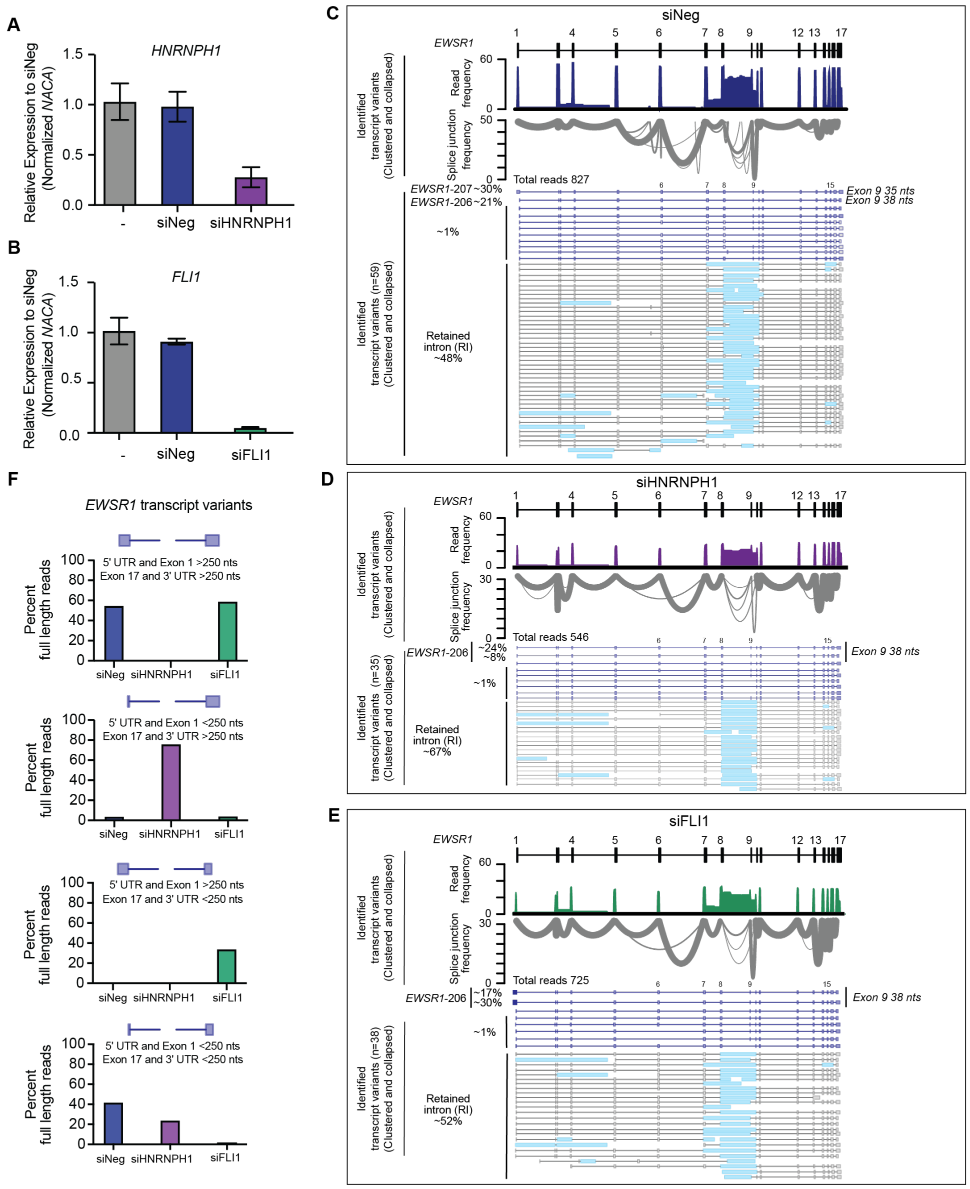


**Figure S1 A**. Quantification of the silencing of *HNRNPH1* in TC32 cells using qPCR analysis (three independent transfections). **B**. Quantification of the silencing of *EWS-FLI1* in TC32 cells using qPCR analysis (three independent transfections). **C**. Gviz views of *EWSR1* transcripts (clustered and collapsed) detected by long-read RNA sequencing of TC-32 cells transfected with the control siRNA, siNeg. **D**. Gviz plots of *EWSR1* transcripts (clustered and collapsed) detected by long-read RNA sequences of analysis of *HNRNPH1*-silenced TC-32 cells. **E.** Gviz plots of *EWSR1* transcripts (clustered and collapsed) detected by long-read RNA sequencing of *EWS-FLI1*-silenced TC-32 cells. **C-E** In each panel, the first track indicates the consensus structure of the *EWSR1* gene. The second track shows the transcript variant mapping frequency. The third track shows the splice junction frequency displayed as Sashimi plots. The fourth track shows schematics of each detected transcript variant (clustered and collapsed). Light blue rectangles indicate retained intronic sequences. The annotated EWSR1 transcripts represent the closest match based on the coding region and its protein-coding potential. **F.** Variations in the length of the 5' or 3' UTRs and the respective adjacent exon (exon 1 or 17) of full-length *EWSR1* transcripts (protein-coding) expressed in TC-32 cells transfected with the indicated siRNAs. We classified the length of each 5' and 3' region (UTR plus the adjacent exon based on a size of greater or less than 250 nts.

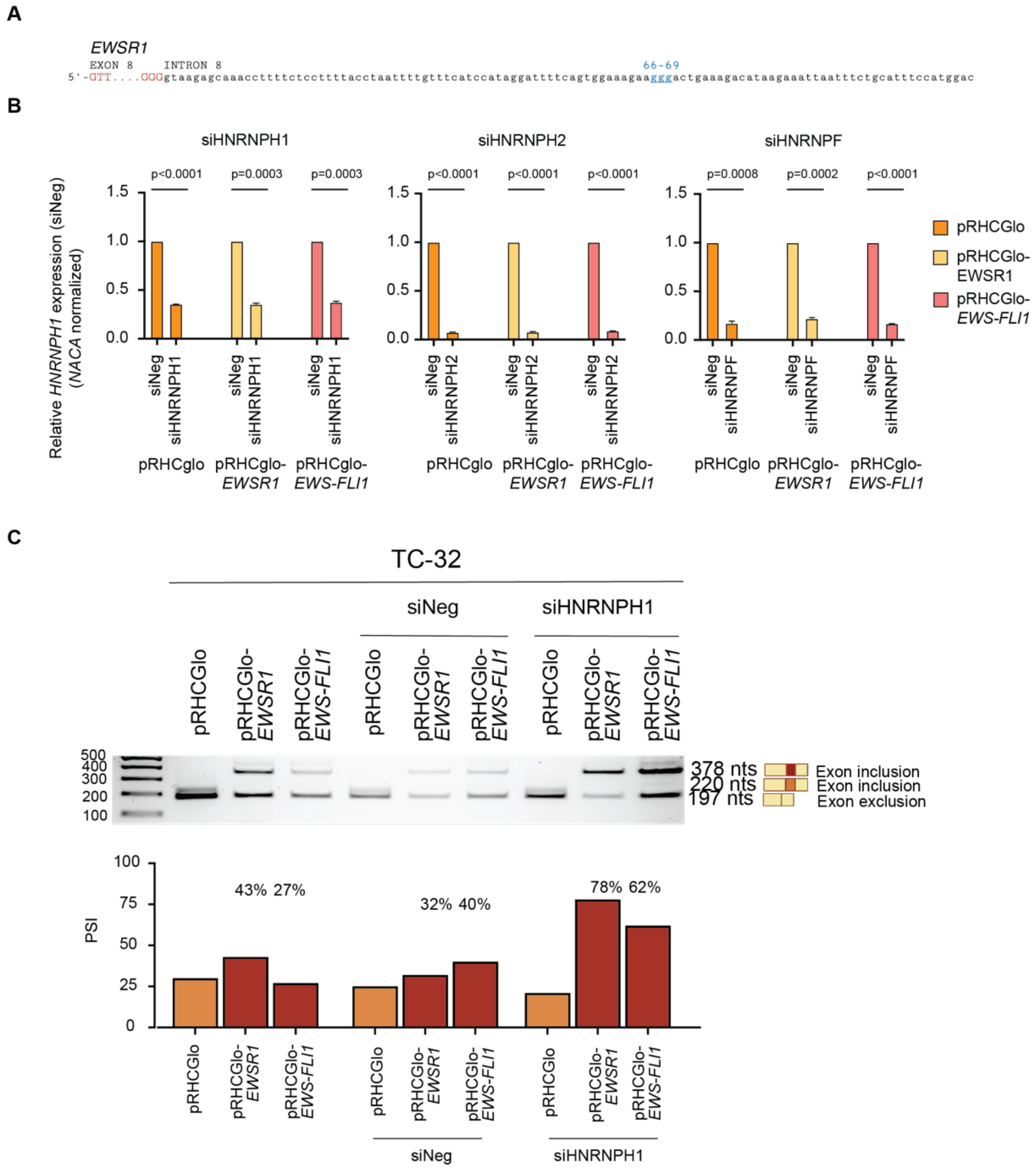


**Figure S2 A.** Schematic of the 5' end of *EWSR1*-intron 8. **B.** Representative qPCR analysis quantifying the silencing of *HNRNPH1*, *HNRNPH2* or *HNRNPF* in HEK-293T cells using the siRNAs targeting each HNRNPH/F family member. We assessed mRNA levels of each transcript species using gene-specific PCR primers, normalized to the expression of NACA and expressed relative to expression in siNeg-transfected cells. The data shown are the mean and standard error of the mean of three transfections. The statistical analysis shows the results of unpaired t tests with Welch's correction. **C.** PCR amplified products obtained following the transfection of TC-32 cells with the indicated plasmids or plasmids and siRNAs, and the quantification of *EWSR1*-exon 8 inclusion (percentage splice inclusion - PSI). The data shown are representative of at least two independent experiments.

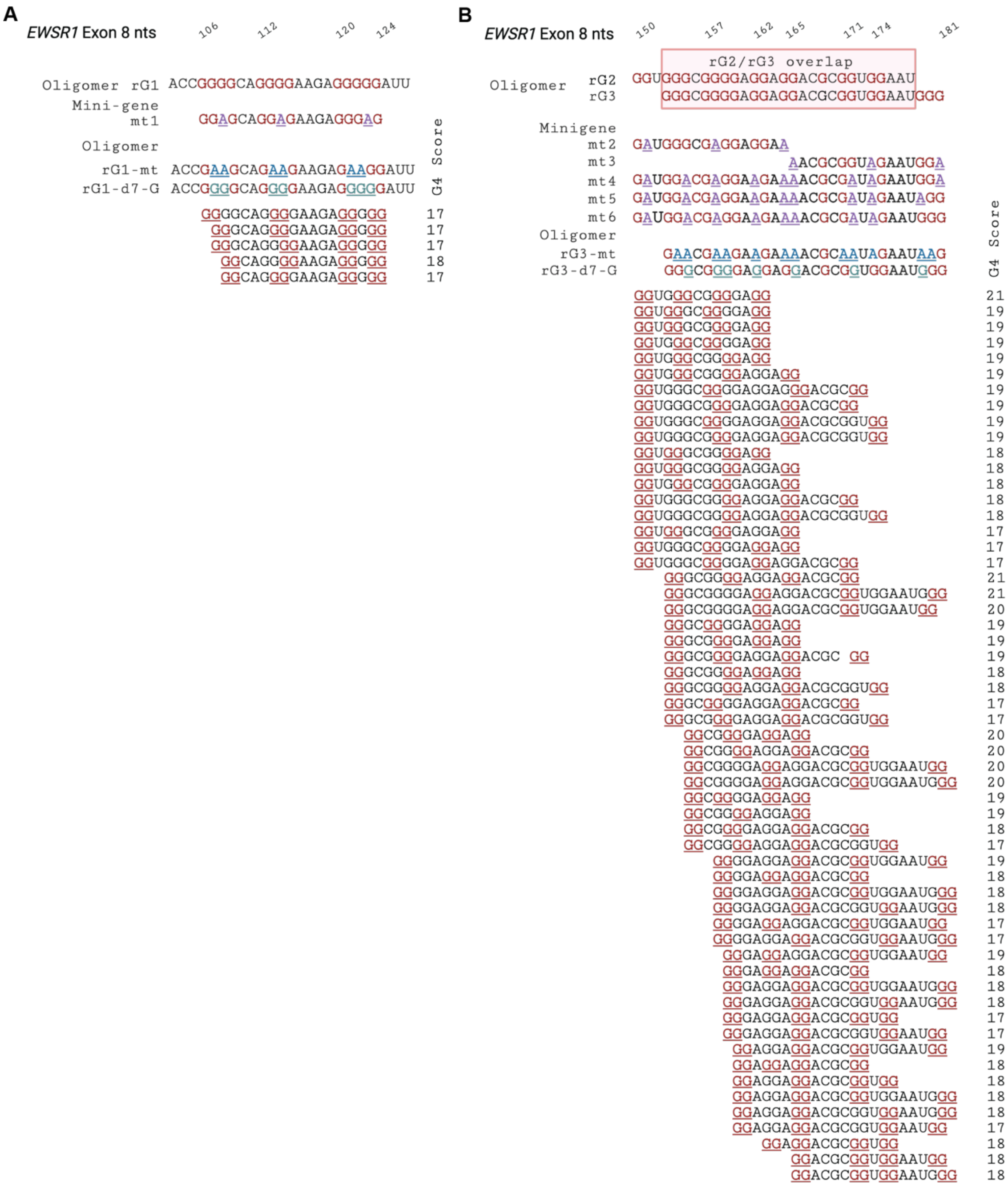

**Figure S3** A summary of guanines that have the potential to become part of two-tier quartet quadruplexes based on the *EWSR1*-exon 8 rG1 and alignment of G-A substitutions used in this study. (**A**), rG1 and (**B**), and rG2/3. Shown are those sequences that the QGRS Mapper (47) predicts form G4 structures (score of 17 or greater) all with a consensus motif of $G_2$-$N_{1\text{-}7}$-$G_2$-$N_{1\text{-}7}$, $G_2$-$N_{1\text{-}7}$-$G_2$. The underlined red Gs indicate those guanines predicted to be part of a quadruplex structure. The underlined blue, green, or purple residues indicate G-A substitutions in the minigenes or G-A and/or d7-G versions of the RNA oligomers used in this study.

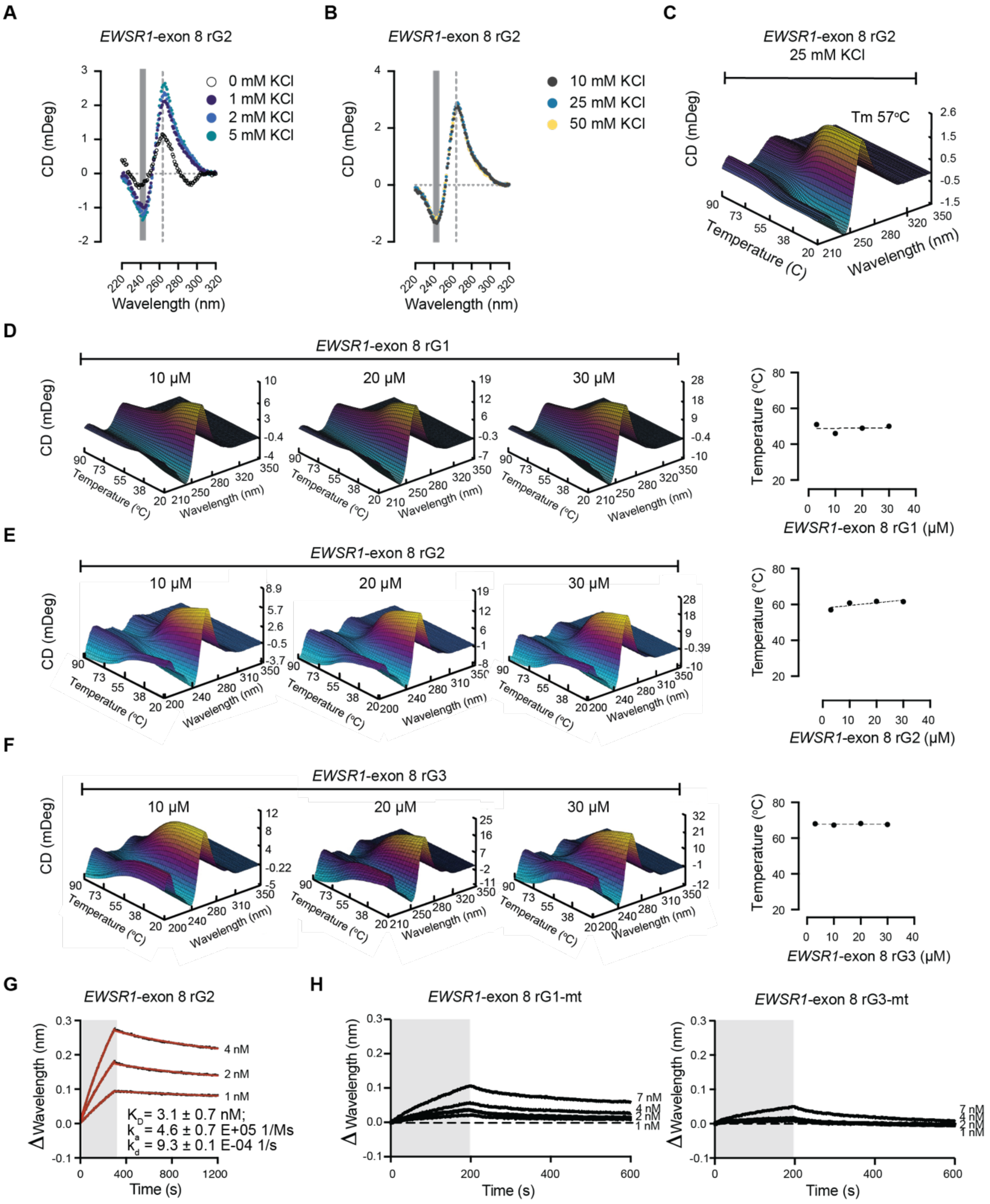


**Figure S4 A.** CD spectra for the rG2 RNA oligomer in the presence of no added KCl, 1, 2, or 5 mM KCl. **B.** CD spectra for the rG2 RNA oligomer in the presence of 10, 25, or 50 mM KCl. **C.** Thermal melt curves for the rG2 RNA oligomer (25 mM KCl). **D.** Thermal melt curves for the rG1 RNA oligomer analyzed at the indicated concentrations and summary of these results. **E.** Thermal melt curves for the rG2 RNA oligomer analyzed at the indicated concentrations and summary of these results. **F.** Thermal melt curves for the rG3 RNA oligomer analyzed at the indicated concentrations and summary of these results. **G.** Sensorgrams for the rG2 RNA oligomer and HNRNPH1 at increasing protein concentrations. **H.** Sensorgrams for the rG1-mt and rG3-mt RNA oligomers and HNRNPH1 at increasing protein concentrations.

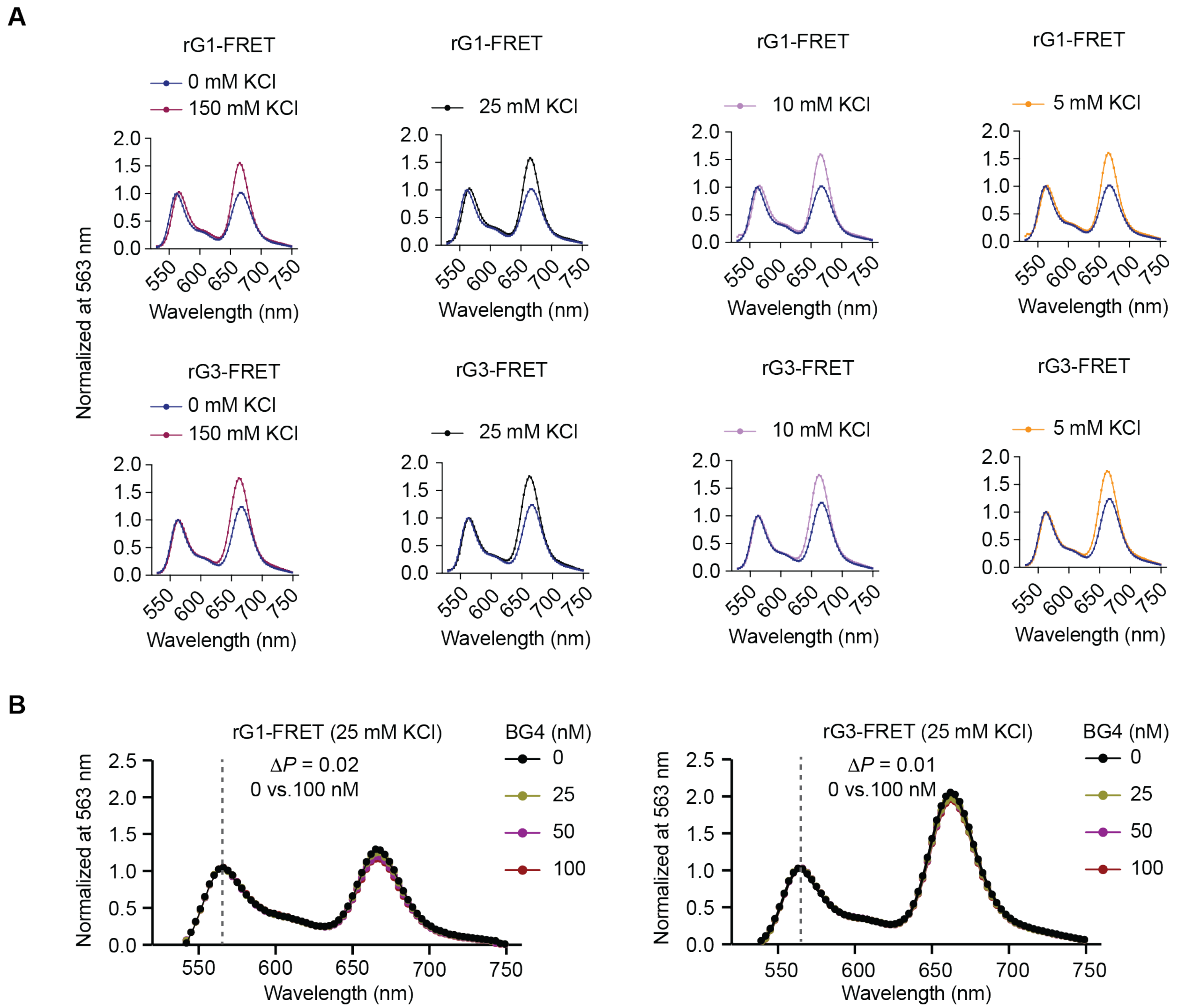


**Figure S5 A.** The individual FRET analysis of the rG1 and rG3 RNA oligomers at the increasing concentrations of KCl shown in **Figure 6C. B.** FRET analysis of the rG1 and rG3 RNA oligomers at increasing concentrations of the pan-G4 antibody, BG4.

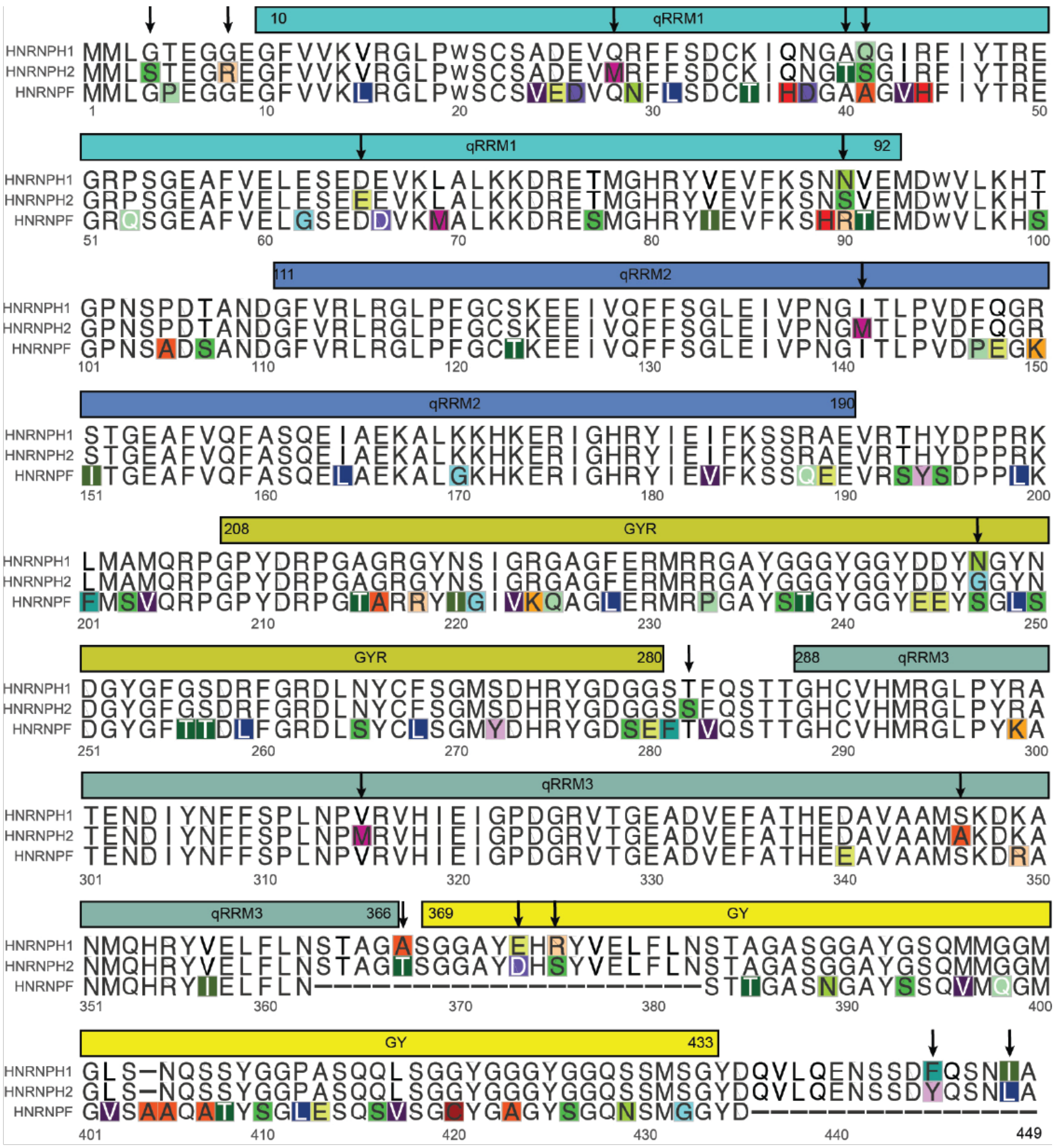

**Figure S6** Alignment of HNRNPH1, HNRNPH2, and HNRNPF indicating the positions of the qRRM and glycine rich domains and those residues that differ between these family members. Alignment and visualization generated using ggmsa (https://github.com/YuLab-SMU/ggmsa).

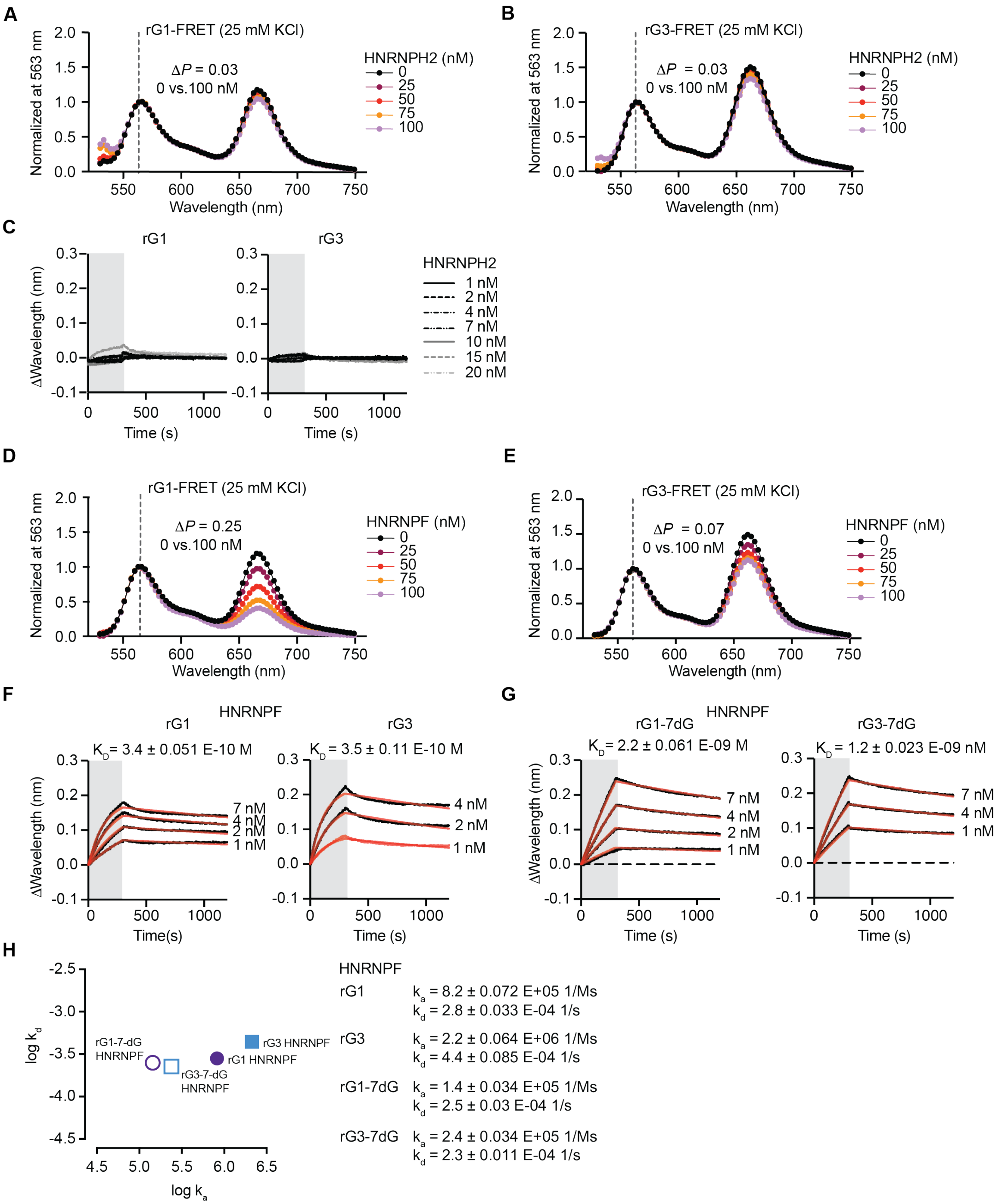


**Figure S7 A.** FRET analysis of the rG1-FRET RNA oligomers at increasing concentrations (0 – 100 nM) of HNRNPH2. **B.** FRET analysis of the rG3-FRET RNA oligomers at increasing concentrations (0 – 100 nM) of HNRNPH2. **C.** Sensograms of the rG1 and rG3 RNA oligomers at increasing protein concentrations of HNRNPH2. **D.** FRET analysis of the rG1-FRET RNA oligomers at increasing concentrations (0 – 100 nM) of HNRNPF. **E.** FRET analysis of the rG3-FRET RNA oligomers at increasing concentrations (0 – 100 nM) of HNRNPF. **F.** Sensograms of the rG1 and rG3 RNA oligomers at increasing protein concentrations of HNRNPF. **G.** Sensograms of the rG1-7-dG and rG3-7-dG RNA oligomers at increasing protein concentrations of HNRNPF. **H.** Graphical representations of the kinetic association and disassociation values of the indicated RNA oligomers in the presence of HNRNPF and a summary of these values.

**A**

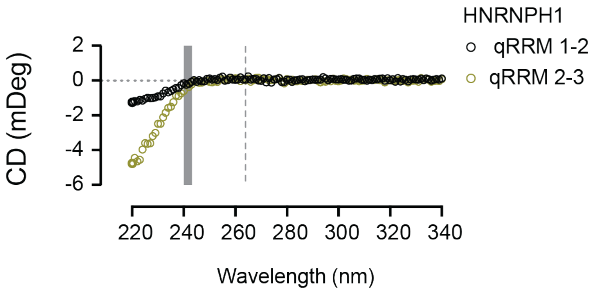


**B**

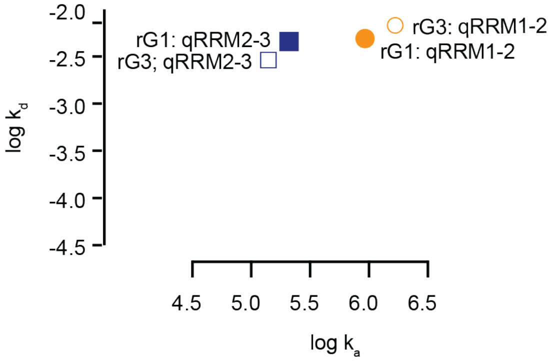


*EWSR1*-exon 8 rG1
HNRNPH1 qRRM1-2
$k_a$ = 9.3 ± 0.1 E+05 (1/Ms)
$k_d$ = 4.9 ± 0.1 E-03 (1/s)

*EWSR1*-exon 8 rG3
HNRNPH1 qRRM1-2
$k_a$ = 16.7 ± 0.04 E+05 (1Ms)
$k_d$ = 6.8 ± 0.09 E-03 (1/s)

*EWSR1*-exon 8 rG1
HNRNPH1 qRRM2-3
$k_a$ = 2.1 ± 0.53 E+05 (1/Ms)
$k_d$ = 4.8 ± 0.04 E-03 (1/s)

*EWSR1*-exon 8 rG3
HNRNPH1 qRRM2-3
$k_a$ = 1.4 ± 0.9 E+05 (1/Ms)
$k_d$ = 2.9 ± 0.9 E-03 (1/s)

**Figure S8 A.** CD spectra for the HNRNPH1 qRRM1-2 and qRRM2-3 domains. **B.** Graphical representations of the association and disassociation values of the indicated RNA oligomers and a summary of these values.

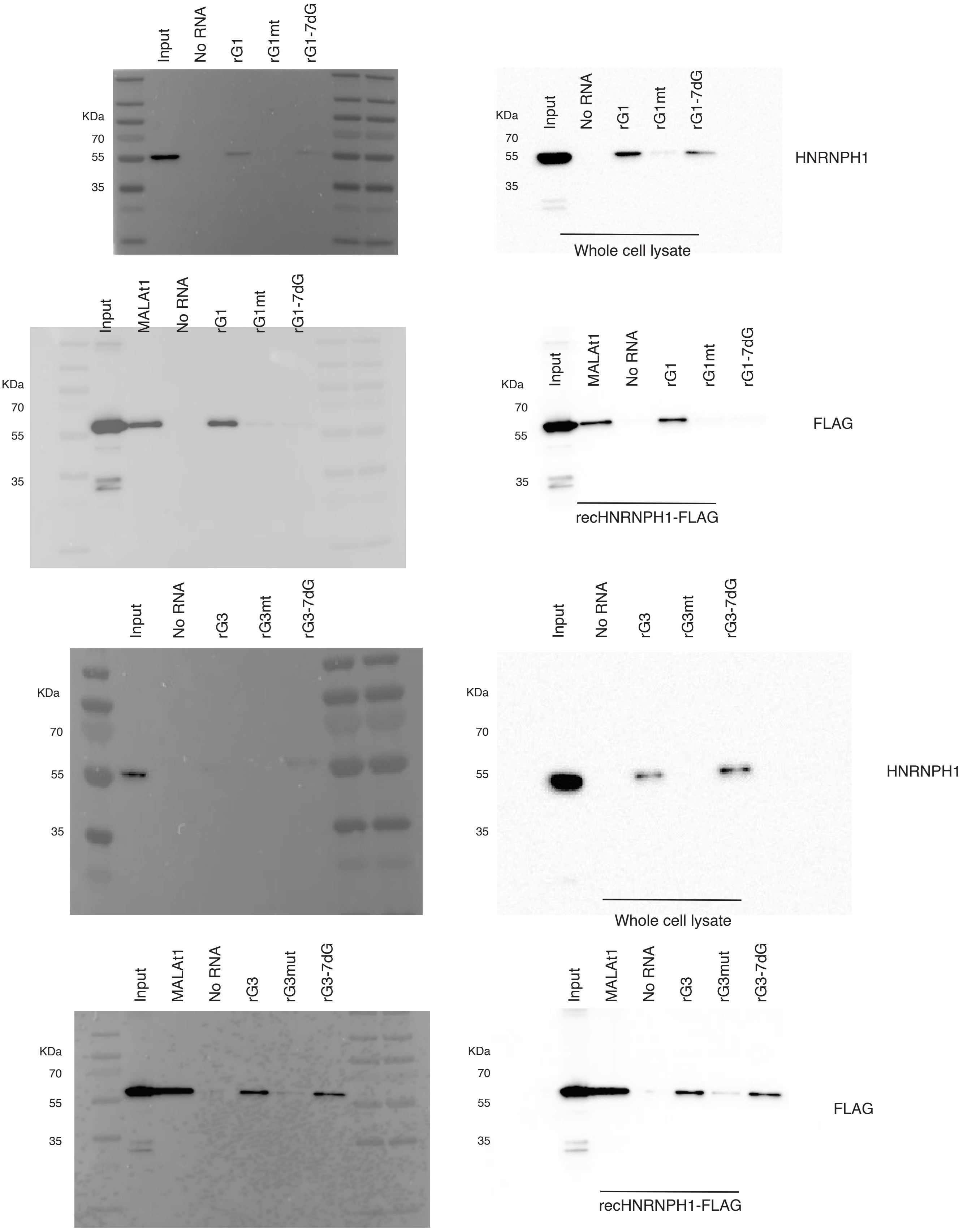
Input
No RNA
rG1
rG1mt
rG1-7dG
KDa
70
55
35
HNRNPH1
Whole cell lysate
MALAt1
FLAG
recHNRNPH1-FLAG
rG3
rG3mt
rG3-7dG
rG3mut
